# A Transcriptional Switch Governing Fibroblast Plasticity Underlies Reversibility of Chronic Heart Disease

**DOI:** 10.1101/2020.07.21.214874

**Authors:** Michael Alexanian, Pawel F. Przytycki, Rudi Micheletti, Arun Padmanabhan, Lin Ye, Joshua G. Travers, Barbara Gonzalez Teran, Qiming Duan, Sanjeev S. Ranade, Franco Felix, Ricardo Linares-Saldana, Yu Huang, Gaia Andreoletti, Jin Yang, Kathryn N. Ivey, Rajan Jain, Timothy A. McKinsey, Michael G. Rosenfeld, Casey Gifford, Katherine S. Pollard, Saptarsi M. Haldar, Deepak Srivastava

## Abstract

In diseased organs, stress-activated signaling cascades alter chromatin, triggering broad shifts in transcription and cell state that exacerbate pathology. Fibroblast activation is a common stress response that worsens lung, liver, kidney and heart disease, yet its mechanistic basis remains poorly understood^1,2^. Pharmacologic inhibition of the BET family of transcriptional coactivators alleviates cardiac dysfunction and associated fibrosis, providing a tool to mechanistically interrogate maladaptive fibroblast states and modulate their plasticity as a potential therapeutic approach^3–8^. Here, we leverage dynamic single cell transcriptomic and epigenomic interrogation of heart tissue with and without BET inhibition to reveal a reversible transcriptional switch underlying stress-induced fibroblast activation. Transcriptomes of resident cardiac fibroblasts demonstrated robust and rapid toggling between the quiescent fibroblast and activated myofibroblast state in a manner that directly correlated with BET inhibitor exposure and cardiac function. Correlation of single cell chromatin accessibility with cardiac function revealed a novel set of reversibly accessible DNA elements that correlated with disease severity. Among the most dynamic elements was an enhancer regulating the transcription factor MEOX1, which was specifically expressed in activated myofibroblasts, occupied putative regulatory elements of a broad fibrotic gene program, and was required for TGFβ-induced myofibroblast activation. CRISPR interference of the most dynamic *cis*-element within the enhancer, marked by nascent transcription, prevented TGFβ-induced activation of *Meox1*. These findings identify MEOX1 as a central regulator of stress-induced myofibroblast activation associated with cardiac dysfunction. The plasticity and specificity of the BET-dependent regulation of MEOX1 in endogenous tissue fibroblasts provides new *trans*- and *cis*- targets for treating fibrotic disease.

---

In many human diseases, dynamic changes in gene expression fuel progressive organ dysfunction. As such, targeting gene transcription has emerged as a new therapeutic strategy in a variety of chronic conditions, including heart failure, a common and lethal condition afflicting 24 million people worldwide^9^. Among strategies to therapeutically target the gene regulatory apparatus, small molecule inhibitors of bromodomain and extra-terminal domain (BET) proteins (BRD2, BRD3, BRD4) have emerged as potent tools to reversibly interdict enhancer-to-promoter signaling *in vivo*^10–12^. BET proteins are a highly conserved family of ubiquitously expressed acetyllysine reader proteins that co-activate transcription^13^, and systemic administration of BET bromodomain inhibitors can ameliorate heart failure in mouse models^3^. Because the cell types most affected by BET inhibition in these models are not known and systemic BET inhibition is too broad to be therapeutically tractable for chronic cardiovascular indications, we leveraged single cell transcriptomic and epigenomic interrogation of heart tissue in the setting of intermittent BET bromodomain inhibitor exposure to reveal cell states and *cis*- and *trans*-targets critical for disease pathogenesis and therapeutic efficacy.

We first investigated whether the therapeutic effect of small molecule BET bromodomain inhibition in mouse models of heart failure was reversible upon compound initiation, withdrawal, and reinitiation. In initial studies, we tested the compound CPI-456 in a mouse model of heart failure induced by a permanent anterior wall myocardial infarction (MI). CPI-456 is an orally bioavailable BET bromodomain inhibitor with subnanomolar potency and drug-like pharmacokinetic properties, initially developed as a clinical candidate for cancer therapy^14^. One month of CPI-456 treatment commenced at post-MI day 5 significantly improved left ventricle (LV) systolic function **(Fig.1A)**. Discontinuation of CPI-456 for the next 3 weeks led to a regression of LV systolic function **(Fig.1A)**. Re-initiation for the next 2 weeks improved LV systolic function to the same degree as the initial treatment phase and once again, subsequent discontinuation of CPI-456 in the final week of the study led to a regression of LV function back to that of untreated controls. We observed similar reversibility of treatment effect using the small-molecule BET-inhibitor JQ1^11^ in a well-established mouse model of LV pressure overload induced heart failure achieved via transverse aortic constriction (TAC) **(Fig.1B)**. Together, these studies using chemically diverse BET bromodomain inhibitors in different murine heart failure models demonstrate significant therapeutic reversibility, with LV function tracking with compound exposure.

**Figure 1:**
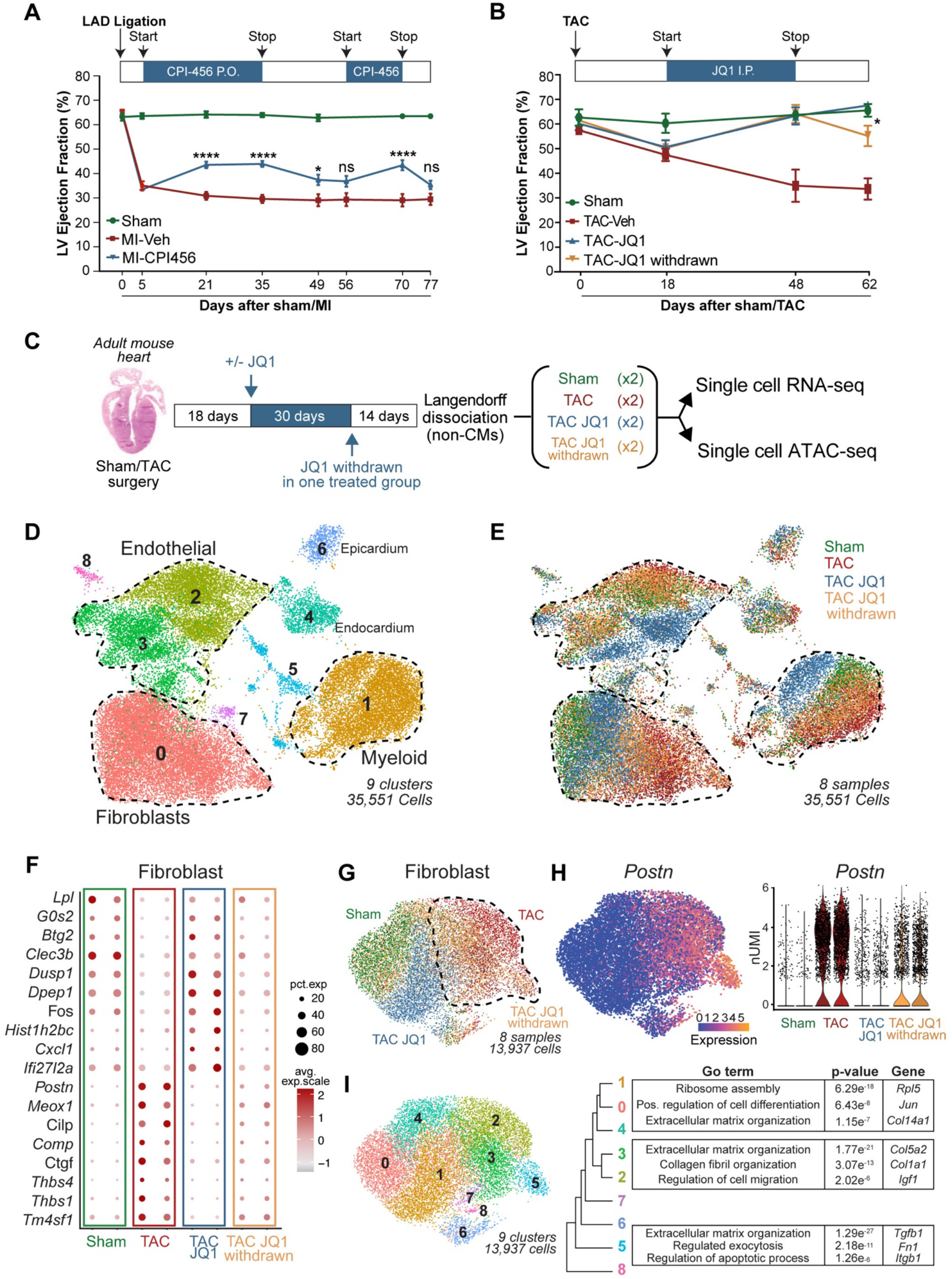
Dynamic reversibility of heart failure with BET bromodomain inhibitor exposure correlates with myofibroblast cell state. **A.** Left ventricle (LV) ejection fraction (EF) quantified by echocardiography (n = 17, 21 and 21 for Sham, MI-Veh and MI-CPI456) in Myocardial infarction (MI) model with intermittent dosing of BET inhibitor CPI456 (10mgk). Statistical significance is shown between MI-Veh and MI-CPI456. **B.** LV EF quantified by echocardiography (n = 4, 6, 6 and 6 for Sham, TAC-Veh, TAC JQ1 and TAC JQ1 withdrawn) in Transverse Constriction Model (TAC) with intermittent dosing of BET inhibitor JQ1 (50mgk). Statistical significance is shown between TAC JQ1 and TAC JQ1 withdrawn. **C.** Experimental workflow for generating single cell RNA- and ATACseq samples from heart samples. **D,E.** UMAP plot of all captured cells in the adult mouse populations colored by cluster (c) and sample (d) identity. Total cells n = 35,551. **F.** Dot plot showing expression (avg.exp.scale) and cell percentage of top differentially expressed (DE) marker genes between samples. **G.** UMAP plot of FB subclustered colored by sample identity. Total cells n = 13,937. **H.** *Periostin* (*Postn*) expression in FBs in the samples shown as UMAP feature plot and violin plot (y axis is normalized UMI levels). **I.** UMAP plot of FBs subclustered colored by cluster identity. Tree diagram showing cluster relationship and representative top Gene Ontology (GO) terms for clusters 0,1,4; 2,3; and 5 are shown to the right. For A. and B., *P < 0.05 and ****P < 0.0001 for indicated comparison. Data are shown as means ± SEM.

As BET bromodomain inhibition reversibly disrupts enhancer-to-promoter signaling^15,16^, we hypothesized that exposure to these compounds could drive reversible changes in cardiac cell states *in vivo* in a manner that correlates with their observed therapeutic efficacy. We performed all subsequent *in vivo* transcriptomic and epigenomic analyses in the mouse TAC model, which exerts stress on all regions of the LV in a stereotypic and highly reproducible manner. Given the striking protective effect of BET inhibition on LV systolic function, we began by profiling cardiomyocytes (CM). As the vast majority of adult CMs are too large to be adequately accommodated in the typical single-cell microfluidic workflow, we isolated adult CMs and analyzed them by bulk RNA-Seq. Surprisingly, we found that the effects of JQ1 on the transcriptome of isolated adult CMs was modest when compared to the previously published transcriptomic signature of whole LV tissue (<3% overlap, **Extended data Fig. 1A**)^5^, strongly suggesting that the most robust effects on gene expression changes were occurring in non-CM populations. Therefore, we performed single cell RNA sequencing (scRNAseq) in the non-CM compartment of mouse hearts using the 10X Genomics platform. We sequenced over 35,000 individual cells collected from four experimental groups: Sham, TAC vehicle-treated (TAC), TAC JQ1-treated (TAC JQ1), and TAC JQ1-treated followed by JQ1 withdrawal (TAC JQ1 withdrawn) **(Fig. 1C)**. Unsupervised clustering of the scRNAseq appropriately identified a diverse array of cardiac cell subpopulations, including FBs, endothelial cells, myeloid cells and epicardial cells **(Fig. 1D and Extended data Fig. 1B,C)**. The most striking finding in this clustering was evident in the FB population, where TAC caused a large shift in cell state and JQ1 treatment lead to a dramatic reversion of this cell state to one that closely approached the Sham state **(Fig. 1E)**. Withdrawal of JQ1 was associated with a shift of the FB population back to a TAC-like stressed state, highlighting a reversible sensitivity of this cellular compartment to JQ1 exposure **(Fig. 1E)**. Interestingly, JQ1 exposure also led to dynamic transcriptomic shifts in the endothelial and myeloid compartments **(Fig. 1E and Extended data Fig. 1D,E)**. However, in contrast to the reversible transitions of FBs between Sham- and TAC-like states, the crisp reversibility of JQ1-mediated shifts in cell state was less evident in endothelial and myeloid cells.

Given the nearly complete bi-directional reversibility of FB cell states in response to JQ1 exposure/withdrawal, we focused our attention on dissecting the transcriptional plasticity of the FB compartment. Differential expression analysis of 13,937 individual FB transcriptomes showed strong similarities between Sham and TAC JQ1 groups and highlighted a core signature of pro-fibrotic genes highly attenuated by BET inhibition **(Fig. 1F)**. Subsetting and reclustering of the FBs further illustrated the robust reversibility of FBs between Sham and TAC states in response to JQ1 exposure **(Fig.1G)**. Cardiac stress is known to trigger the transition of resident FBs into a contractile and synthetic state called the myofibroblast (myoFB)^17,18^. Overlay of the myoFB marker gene *Postrn*^19^ demonstrated that TAC leads to myoFB activation, administration of JQ1 shifts the myoFB cell state back toward a Sham-like state, and withdrawal of JQ1 reverts these cells back to myoFBs **(Fig.1H)**. Sub-clustering the FBs defined 10 clusters with demarcation of basal FB states (encompassing Sham and TAC JQ1 cells; clusters 0, 1, and 4) versus the myoFB state (encompassing TAC and TAC JQ1 withdrawn cells; clusters 2, 3, and 5) **(Fig.1I and Extended data Fig. 1F)**. Gene ontology (GO) analysis highlighted how gene-programs associated with basal FB homeostasis were enriched in Sham and TAC JQ1 cells, while the TAC and TAC JQ1 withdrawn populations were enriched for pro-fibrotic, secretory, proliferative and migratory gene programs **(Fig.1I)**. Together, these data demonstrate that the reversible transition between the basal FB and activated myoFB states can be robustly toggled using transcriptional inhibition, suggesting that BET protein function in myoFBs influence the trajectory of heart failure pathogenesis. Given the dynamic regulation of fibrosis-inducing and secretory proteins in FBs the effects of BET inhibition may be cell autonomous and non-cell autonomous.

Since BET proteins have the potential to regulate chromatin status^15,16^, we hypothesized that the observed transcriptional reversibility that results from JQ1 exposure would be supported by corresponding changes in chromatin accessibility and enhancer activation in cardiac FBs and other endogenous cardiac cell types during heart failure pathogenesis. To test this, we integrated the single cell transcriptomic analysis with single cell Assay for Transposase-Accessible Chromatin sequencing (scATACseq) from the same hearts used for scRNAseq **(Fig. 1C and Extended data Fig. 2A)**^20,21^. We identified 490,020 accessible sites distributed among 31,766 individual cells and assigned cellular identity based on chromatin signature **(Extended data Fig. 2B, C)**. As the focus of this study was to dissect distal regulatory elements, we excluded accessible sites in promoters and gene bodies and defined a catalog of FB-, myeloid- and endothelial-enriched distal elements that were used for all subsequent analyses **(Extended data Fig. 2D)**. Interestingly, FBs showed a significantly greater increase in chromatin accessibility after TAC that was reversibly attenuated with JQ1 treatment **(Fig. 2A)**, a feature that was less evident in myeloid and endothelial cells **(Extended data Fig. 2E, F)**. This highlights how the FB cell population preferentially undergoes chromatin activation during chronic heart failure that is partially dependent on BET proteins. In order to dissect dynamic and reversible changes in chromatin activation, we defined open and closed distal elements across our 4 samples and excluded the regions that were constitutively open across all conditions **(Extended data Fig. 2G)**. We observed robust reversibility of chromatin states in response to stress and BET inhibition, particularly in FBs **(Fig. 2B and Extended data Fig. 2G)**. We discovered a cluster of very sensitive and highly dynamic FB distal elements that were closed in Sham, opened in TAC, closed by JQ1, and robustly re-accessible following JQ1 withdrawal **(Cluster 2, Fig. 2B)**. GO analysis showed that these regions were in proximity of genes controlling heart growth and extracellular matrix (ECM) organization, two hallmark features of adverse cardiac remodeling and fibrosis. Interestingly, we also identified a large cluster of FB regions that opened from Sham to TAC which were insensitive to JQ1, highlighting a signature of stress-responsive chromatin activation that is BET-independent **(Cluster 9, Fig. 2B)**. We next explored how transcription factor (TF) binding motif accessibility changed in regions that were dynamically modulated in the 3 phenotypic transitions where significant changes in heart function occur: Sham to TAC, TAC to TAC-JQ1 and TAC-JQ1 to TAC-JQ1 withdrawn. In FBs, TF binding motifs for CEBPB, JUN and MEOX1^22^ showed enrichment in accessible regions in the Sham to TAC transition and subsequent loss of enrichment with BET inhibition that was then re-acquired with JQ1 withdrawal, suggesting that chromatin dynamics occur at regions enriched with functionally relevant motifs for stress-activated TFs **(Fig. 2C and Extended data Fig. 3A, B)**.

**Figure 2:**
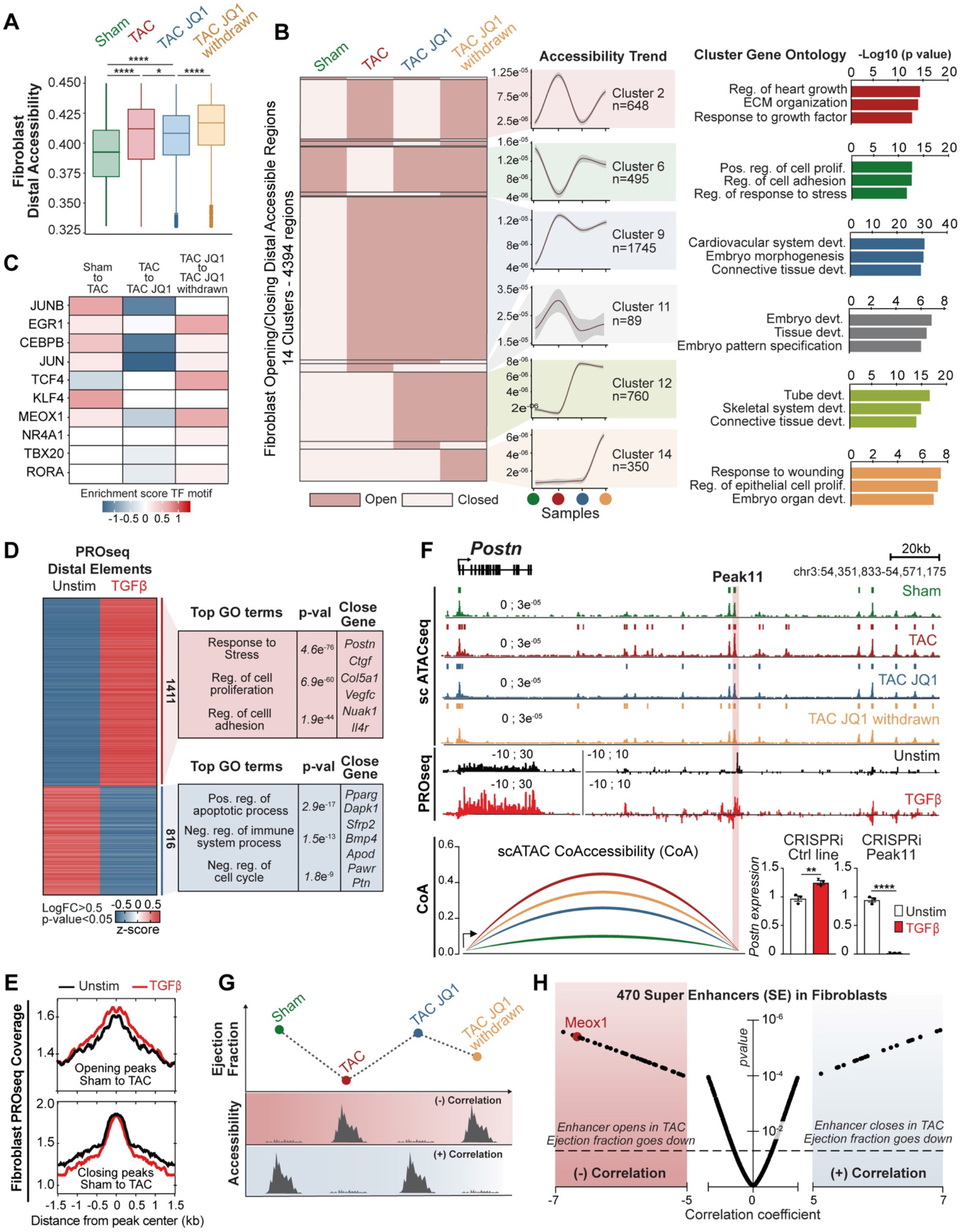
Reversibility of fibroblast chromatin states reveals novel dynamically accessible DNA elements that correlate with heart function. **A.** Chromatin accessibility of distal elements in FB cells derived from scATAC-seq samples. Trimming of 10% most extreme points was performed for better visualization. **B.** Dynamic accessibility of distal elements in FBs clustered by trend across samples (left) with top three GO terms for nearest genes to distal elements in each cluster (right). **C.** Enrichment scores for TF motif accessibility in distal elements between samples for the ten most expressed TFs in TAC in FBs. **D.** Heatmap of PROSeq coverage of differentially transcribed distal regions between Unstimulated (Unstim) and TGFβ-treated FBs. TOP GO terms are shown to the right. Average signal for 2 replicates each condition is shown. **E.** PROseq coverage measured in Unstim and TGFβ-treated FBs *in vitro* of scATAC peaks opening (n=8964) or closing (n=1628) between Sham and TAC *in vivo.* Top related GO terms and example genes are shown to the right. **F.** Coverage of scATAC in FBs and PROseq at the *Postn* locus. A strongly transcribed region (Peak11) is highlighted in red within the large *Postn* enhancer. CRISPRi targeting Peak11 is shown on the bottom right. *Postn* expression by qPCR between Unstim and TGFβ-treated FBs in CRISPRi control line and Peak11 targeting (each panel is normalized to its Unstim condition). **G.** Schematic of correlation analysis between LV ejection fraction and chromatin accessibility highlighting a negative or positive correlation. **H.** Volcano plot showing correlation coefficients (referred to analysis depicted in Fig. 2G) and corresponding *p*-values of 470 super enhancers in FBs. A region distal to the *Meox1* gene has one of the most negative correlation coefficients. For F., **P < 0.01 and ****P < 0.0001 for indicated comparison. Data are shown as means ± SEM.

We next sought to determine functionally relevant FB and activated myoFB enhancers discovered during scATACseq. A body of recent work has shown that enhancers can be pervasively transcribed and that this nascent transcriptional activity is a robust and independent indicator of enhancer activity^23^. Guided by this rationale, we performed precision nuclear run-on sequencing (PROseq)^24^ on cultured FBs *in vitro* to map genome-wide RNA polymerase II nascent transcription and identify putatively active enhancers. Given PROseq requires large quantities of cells, we generated an immortalized line derived from primary adult mouse cardiac FBs and treated it with TGF-β, a canonical stimulus for eliciting myoFB cell state transition *in vitro* **(Extended data Fig. 3C, D)**^25^. We identified a set of distal elements that were significantly more transcribed after TGF-β stimulation and located close to pro-fibrotic and pro-synthetic genes **(Fig. 2D)**. Using our scATACseq data, we identified the distal elements that were either opening or closing between Sham and TAC *in vivo,* and assessed the PROseq signal in the cultured FBs at these same regions. We found that *in vitro* TGF-β stimulation of cultured FBs triggers global transcriptional changes that resemble those that occur in endogenous FBs *in vivo* during heart failure pathogenesis **(Fig. 2E)**. Visualization of the *Postn* locus illustrates this dynamic regulation **(Fig. 2F),** where there is chromatin opening *in vivo* after TAC and dynamic sensitivity to JQ1 exposure. Within this large enhancer, PROseq revealed a specific region that was heavily transcribed following TGF-β treatment (Peak11) **(Fig. 2F)**. scATACseq co-accessibility analysis between the *Postn* promoter and the Peak 11 region showed low co-accessibility in the Sham state, a robust increase in co-accessibility in response to TAC, and modulation of co-accessibility in response to JQ1 exposure **(Fig. 2F)**. We used CRISPR interference (CRISPRi) deploying a catalytically inactive Cas9 protein (dCas9) fused to the KRAB repressor protein^26^ with a guide strand specific to the Peak11 region to drive sequence-specific repression of this particular regulatory element in FBs, and found that this region is essential for *Postn* induction following TGF-β stimulation **(Fig. 2F and Extended data Fig. 3E)**.

Having demonstrated the dynamic transcriptional control of *Postn,* a known marker of myoFBs^19^, we hypothesized that our integrated single cell transcriptomic and epigenomic approach could be leveraged to discover novel mechanisms controlling cellular stress responses during disease pathogenesis. We built an unbiased enhancer discovery pipeline to unveil distal elements that could play a role in the progression and reversal of heart failure. We assembled a catalog of cell population-enriched large enhancers (also known as stretch- or super-enhancers)^27,28^ in FBs, myeloid and endothelial cells using our scATACseq data in the diseased heart (TAC) **(Extended data Fig. 3F)**. As BET inhibition robustly improved heart function in the TAC model, we correlated the degree of accessibility of these enhancers in FB, myeloid and endothelial cells with LV ejection fraction. This correlation analysis between a measure of enhancer chromatin accessibility and a physiological trait (in this case LV ejection fraction) is summarized in **Figure 2G**. Enhancer elements were defined as having a negative correlation if their accessibility was anti-correlated with heart function (i.e., these enhancers were opening from Sham to TAC, a setting where cardiac function decreases). Conversely, enhancers with a positive correlation were those that closed from Sham to TAC. A Volcano plot of correlation coefficient was generated for each cell type **(Fig. 2H and Extended data Fig. 3G, H)**. Of the 470 large enhancers identified in FBs, 48 showed strong negative correlation while 22 showed strong positive correlation **(Fig. 2H)**.

Notably, one of the most negatively correlated elements in FBs was a large enhancer downstream of *Meox1* **(Fig. 2H and Extended data Fig. 4A)**, a homeodomain-containing TF that is expressed in paraxial mesoderm and is required for sclerotome development^29,30^. *Meox1* was particularly interesting because it was minimally expressed in the healthy mouse heart but highly upregulated in MyoFBs following TAC **(Fig. 3A and Extended data Fig. 4B)**. BET inhibition abolished *Meox1* expression while JQ1 withdrawal was associated with its robust re-induction **(Fig. 3A)**. This, combined with the corresponding enrichment of the MEOX1 DNA-binding motif in dynamically accessible regions of chromatin in endogenous cardiac FBs **(Fig.2C)**, suggested dynamic upregulation and functional engagement of MEOX1 with key FB regulatory elements under stress conditions.

**Figure 3:**
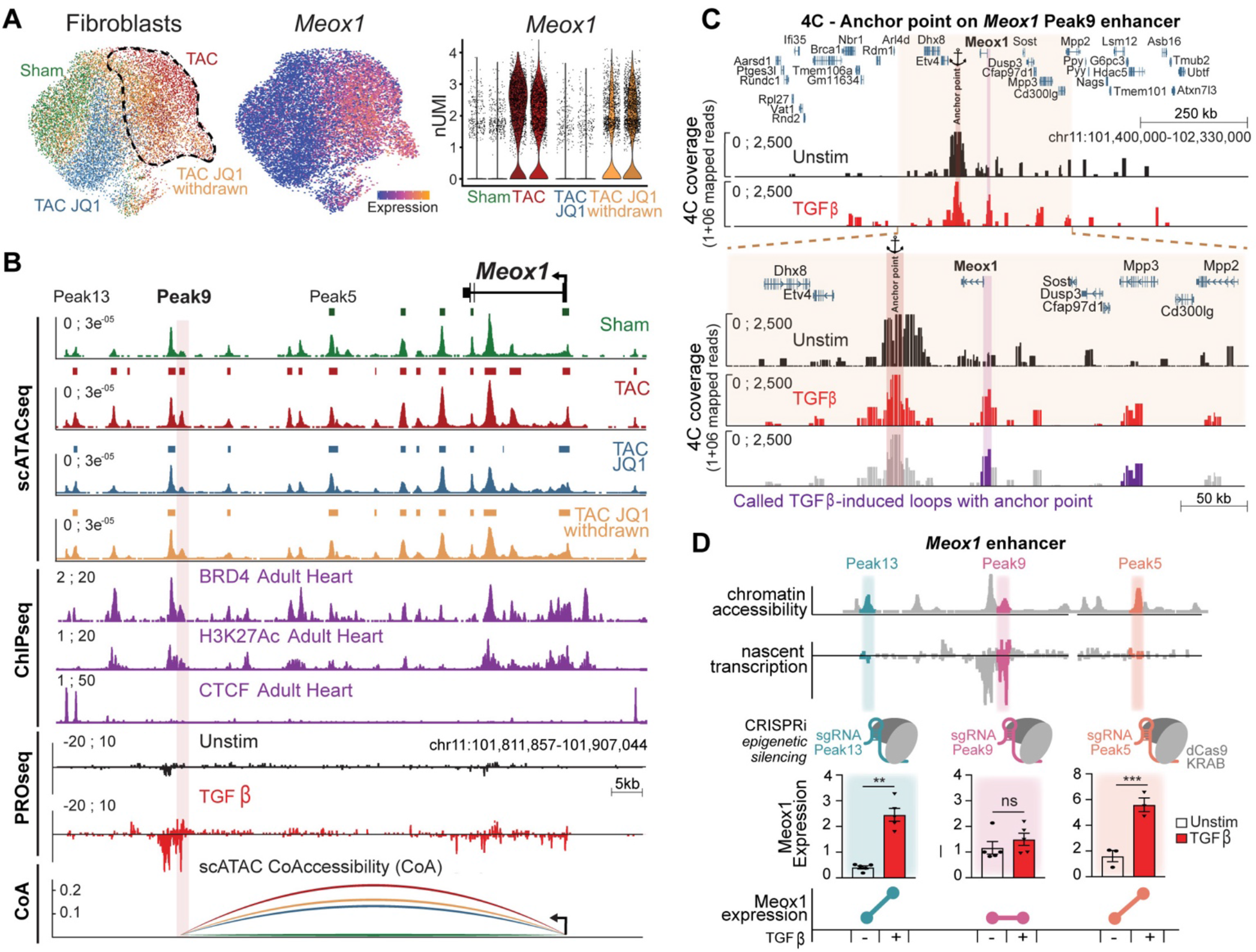
Chromatin accessibility and nascent transcription identify a cis-regulatory element controlling Meox1 expression. **A.** UMAP plot of FBs subclustered colored by sample identity (same as Fig.1G) and *Meox1* expression in FB in the samples shown as UMAP feature plot and violin plot (y axis is normalized UMI levels). **B.** *Meox1* locus (gene and enhancer) showing from top to bottom: coverage of scATAC samples in FBs; ChIPseq for BRD4 (GSE46668), H3K27Ac and CTCF (ENCSR000CDF and ENCSR000CBI) in the adult heart; coverage of PROseq in Unstim and TGFβ-treated FBs; and co-accessibility measures between Meox1 promoter and Peak9 region in FBs using scATAC. A highly transcribed region (Peak9) is highlighted in red within the large *Meox1* enhancer. **C.** Chromosome conformation capture (4C) between the Peak9 region (anchor point) and *Meox1* promoter showing 4C coverage in Unstim and TGFβ-treated FBs. 922kb (top) and 328kb (bottom) genomic regions are shown. Last track represents the called TGFβ-induced loops with Peak9 (colored in purple). **D.** Schematic showing CRISPRi targeting of 3 regions within the *Meox1* enhancer (Peak5, 9 and 13) at the top. *Meox1* expression by qPCR between Unstim and TGFβ-treated FBs in the 3 CRISPRi FB lines targeting Peak5, 9 or 13 (each panel is normalized to its Unstim condition). For D., **P < 0.01 and ***P < 0.001 for indicated comparison. Data are shown as means ± SEM.

The enhancer downstream of *Meox1* was extremely sensitive to stress and JQ1 exposure in FBs, but not in myeloid and endothelial cells **(Fig. 3B and Extended data Fig. 4C)**. The enhancer significantly opened from Sham to TAC conditions in FBs with 10 peaks that became accessible in this transition, closed with JQ1 treatment back to a Sham level, and re-opened when JQ1 was withdrawn **(Fig. 3B and Extended data Fig. 4C,D)**. Analysis of the chromatin accessibility in all individual peaks of the *Meox1* enhancer showed how particular elements were dynamically modulated **(Extended data Fig. 4E)**. Publicly available BRD4 and H3K27ac ChIP-Seq data from adult mouse LV tissue corroborated these active enhancer marks at the *Meox1* locus, and CTCF ChIPseq was consistent with the absence of contact insulation between the *Meox1* gene and the enhancer, raising the possibility that this enhancer regulates *Meox1* **(Fig. 3B)**.

Importantly, *Meox1* mRNA expression was induced in cultured FBs treated with TGF-β **(Extended data Fig. 5A)**. TGF-β-induced *Meox1* upregulation was suppressed by JQ1 and knockdown of each of the 3 BETs individually with siRNAs demonstrated that *Meox1* induction was dependent on BRD4, but not BRD2 or BRD3 **(Extended data Fig. 5A-C)**. Among the individual scATAC-Seq peaks in this locus that showed increased accessibility with TAC *in vivo,* PROseq in cultured FBs was able to identify a specific region located 62 kilobases (kb) downstream of the *Meox1* promoter (Peak 9) that featured a striking increase in nascent transcription following TGF-β stimulation **(Fig. 3B)**. Notably, this 780 base pair (bp) element showed stronger TGF-β stimulated transcription than the *Meox1* gene body itself and was also one of the most differentially transcribed regions across the whole genome in response to TGF-β stimulation **(Fig. 3B and Extended data Fig. 6A)**. The *Meox1* promoter and the Peak 9 region showed low co-accessibility in the Sham state, a strong increase in co-accessibility in response to TAC, and modulation of co-accessibility in response to BET inhibition **(Fig. 3B)**. Chromosome conformation capture analysis of this locus in cultured FBs revealed a robust increase in contact between the Peak 9 enhancer region and the *Meox1* promoter in response to TGF-β stimulation **(Fig. 3C and Extended data Fig. 6B)**, consistent with dynamic contact between these elements. Compared to the other regions within the large *Meox1* regulatory element, Peak 9 featured strong chromatin accessibility and nascent transcription, two features that are strong predictors of a functionally relevant enhancer^23^. To definitively interrogate the endogenous function of Peak 9, we performed a series of CRISPRi experiments in the *Meox1* locus using guide strands specifically targeted to individual sites and found that the Peak 9 element was required for *Meox1* transactivation upon TGF-β stimulation while other accessible regions identified *in vivo* were not **(Fig. 3D and Extended data Fig. 6C,D)**.

We next investigated the function of MEOX1, hypothesizing that this poorly characterized homeobox TF might directly regulate gene programs involved in fibrotic disease. Knockdown of *Meox1* by a siRNA led to significant reduction in TGF-β-stimulated collagen-gel contraction and EdU incorporation, confirming that MEOX1 was required for contractile and proliferative phenotypic transitions, two functional hallmarks of MyoFBs in disease pathogenesis **(Fig. 4A-C and Extended data Fig. 7A,B)**. ChIPseq showed that MEOX1 binds genes involved in FB homeostasis and response to stress **(Fig. 4D)**. GO analysis of the highly MEOX1-bound genes showed enrichment for terms linked to apoptosis, ECM/collagen organization and cell adhesion. In order to understand whether MEOX1 controls the transcription of stress-responsive pro-fibrotic genes, we performed PROseq in TGF-β treated FBs in the presence of either a control or a *Meox1*-targeting siRNA. 509 genes were significantly less transcribed when *Meox1* was depleted, while 819 were more transcribed **(Fig. 4E and Extended data Fig. 7C,D)**. Notably, GO analysis of the *Meox1*-dependent genes revealed enrichment for pro-fibrotic processes such as regulation of cell motility, proliferation, and migration. Among these genes were classical markers of cardiac MyoFB activation, including *Ctgf* and *Postn,* which showed MEOX1 enrichment at their promoters and proximal regulatory elements (including the *Postn* Peak11 enhancer described in **Figure 2F**), regions that featured strong decrease in transcription following *Meox1* depletion **(Fig. 4F)**. These findings suggest MEOX1 functions as an essential transcriptional mediator of the FB to myoFB switch associated with fibrotic disease. Finally, using recently publicly available single cell data from the human adult heart (www.heartcellatlas.org)^29^, we found that *MEOX1* was specifically expressed together with *POSTN* in a subset of activated FBs **(Fig. 4G)**. Notably, *MEOX1* expression was significantly up-regulated in human diseases that prominently feature fibrosis, such as heart tissue from patients with cardiomyopathy and lung tissue from patients with idiopathic pulmonary fibrosis **(Fig. 4H,I)**^30^.

**Figure 4:**
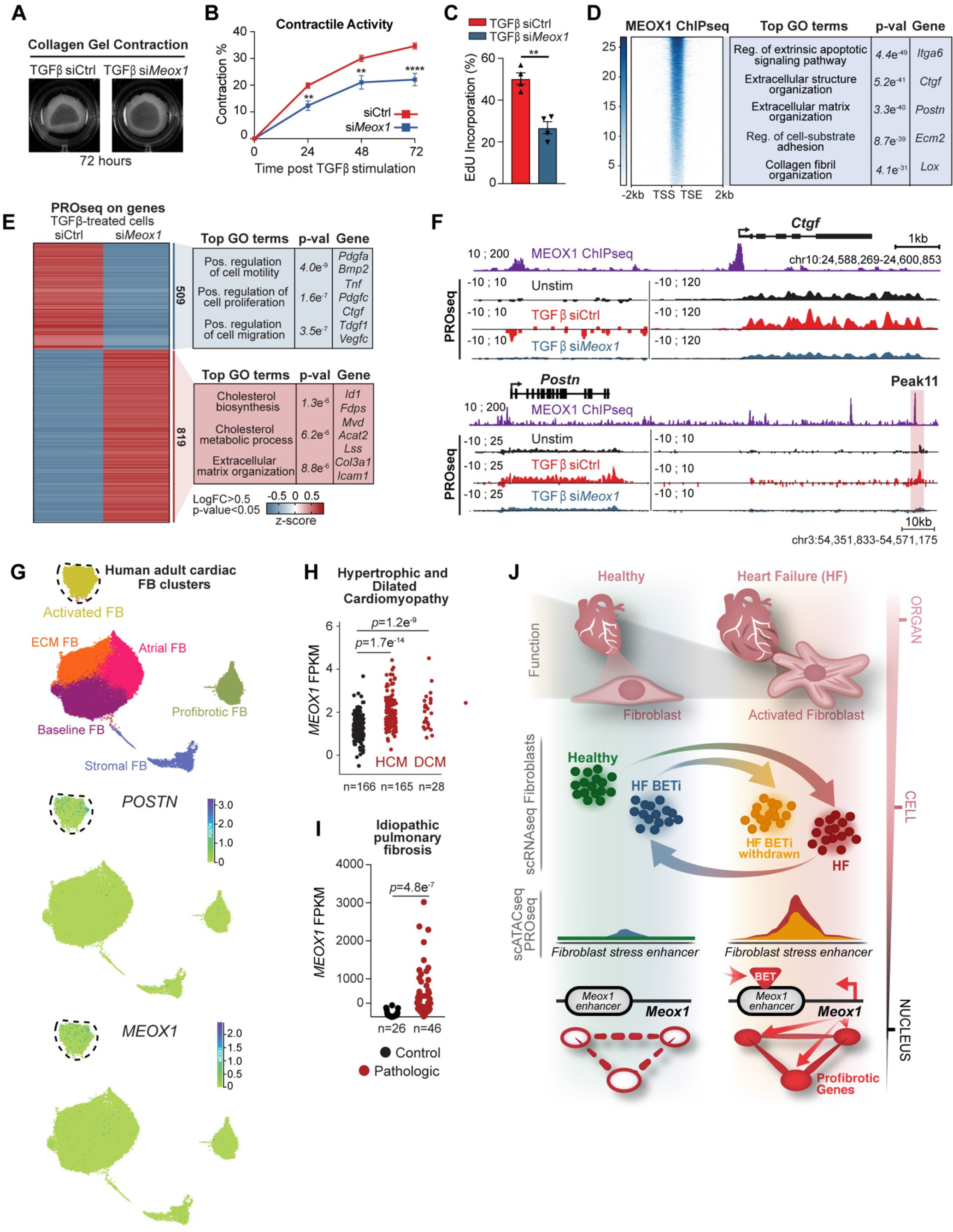
MEOX1 is a novel regulator of fibroblast plasticity and profibrotic function. **A.** Representative images of FBs seeded on compressible collagen gel matrices assayed for gel contraction after treatment with TGFβ and a control (Ctrl) siRNA or one targeting *Meox1* for 72 h. **B.** Quantification of gel contraction images, reported as percentage contraction; (n=4 plates per condition). Data are presented as mean±SEM. **C.** Quantification Edu incorporation in FBs after treatment with TGFβ and a Ctrl or *Meox1* siRNA for 72 h. Data are presented as mean ± SEM. **D.** Heatmap of MEOX1-HA ChIPseq occupancy at protein coding gene (−2kb from transcriptional start site, +2kb from transcriptional end site) sorted by the strength of ChIPseq signal (8366 regions shown). **E.** Heatmap of PROseq coverage of differentially transcribed protein coding genes between TGFβ-treated FBs with Ctrl or *Meox1* siRNA. Average signal for 2 replicates in each condition is shown. Top related GO terms and example genes are shown to the right. **F.** Coverage of MEOX1 ChIP and PROseq (Unstim and TGFβ-treated FBs with Ctrl or Meox1 siRNA) at the *Ctgf* or *Postn* locus (including the *Postn* Peak11 regulatory element) **G.** UMAP plot of adult human cardiac FBs colored by cluster identity (https://www.heartcellatlas.org/fibroblasts). Expression of *POSTN* and *MEOX1* are shown as UMAP feature plots to the right. **H,I.** Bulk RNAseq data of human *MEOX1* expression between controls and individuals with HCM/DCM (h - GSE141910) or Idiopathic pulmonary fibrosis (i - GSE134692), assessed in heart or lung tissue, respectively. **J.** The transcriptional switch that activates fibroblasts correlates with heart disease state. Combining single cell transcriptomic and epigenomic interrogation, we discovered key enhancers and protein coding genes that dynamically regulate fibroblast plasticity and profibrotic function, including the transcription factor MEOX1. For B. and C., **P < 0.01 and ****P < 0.0001 for indicated comparison. Data are shown as means ± SEM.

This study uncovers the transcriptional switch for cellular plasticity in the FB compartment during chronic heart failure pathogenesis and demonstrates that this maladaptive cell state transition is a druggable feature of disease. Using single cell transcriptomic and epigenomic interrogation of failing mouse hearts, we have discovered key enhancers and protein coding genes that dynamically regulate FB cell state, exemplified by the identification of the homeodomain transcription factor MEOX1 as a novel regulator of FB plasticity and profibrotic function **(Fig. 4J)**. This work highlights that single-cell based interrogation of cell states in a diseased tissue, coupled with temporally-controlled perturbation of transcription signaling, can be leveraged to discover the plasticity of cell states and molecular mechanisms critical for progression and reversal of fibrotic diseases. Mechanistic refinement converging on cell-type specific enhancers in the context of a complex tissue offers the opportunity to develop therapeutic approaches that are tailored to targeted gene regulation in defined cell compartments, in contrast to the broad effects of BET inhibition. These findings may inform new therapeutic strategies for a wide variety of chronic disorders that feature maladaptive remodeling of cell state and tissue architecture.

## Acknowledgments

We thank the Srivastava laboratory for critical discussions and feedback; Joke van Bemmel, Mauro Costa, Nathan Palpant, Phillip Grote, Elphège P. Nora, Arjun A. Rao and Alexis J. Combes for the helpful discussions; Jun Qi and Deyao Li for kindly providing the JQ1 molecule; we acknowledge the Gladstone Genomics Core and Flow Cytometry Core for their technical expertise and the Gladstone Animal Facility for support with mouse colony maintenance; and Ana Catarina Silva for helping with figure editing and design.

## Author contributions

M.A., S.M.H and D.S. conceived the study, interpreted the data and wrote the manuscript. P.F.P and K.S.P analyzed scATAC data. R.M. and M.G.R performed and analyzed PROseq and 4C. M.A and A.P. isolated cells/nuclei for scRNA/scATACseq. M.A, L.Y and F.F. performed TGFβ stimulation, CRISPRi assays and generated Meox1-HA line. J.G.T and T.A.M performed collagen contraction and EdU incorporation assays. M.A. and B.G.T generated immortalized fibroblast line and performed ChIPseq. M.A. and Q.D. performed langendorff perfusion. S.S.R. prepared scRNA/ATAC libraries. M.A, R.L.S and R.J analyzed ChIPseq data. Y.H. and J.Y. performed TAC/MI surgeries and analyzed echocardiography data. M.A. and G.A. analyzed bulk RNAseq. K.N.I supervised CPI-456 study. M.A. and C.G. performed scRNAseq.

## Extended Data Figure

**Extended data Figure 1:**
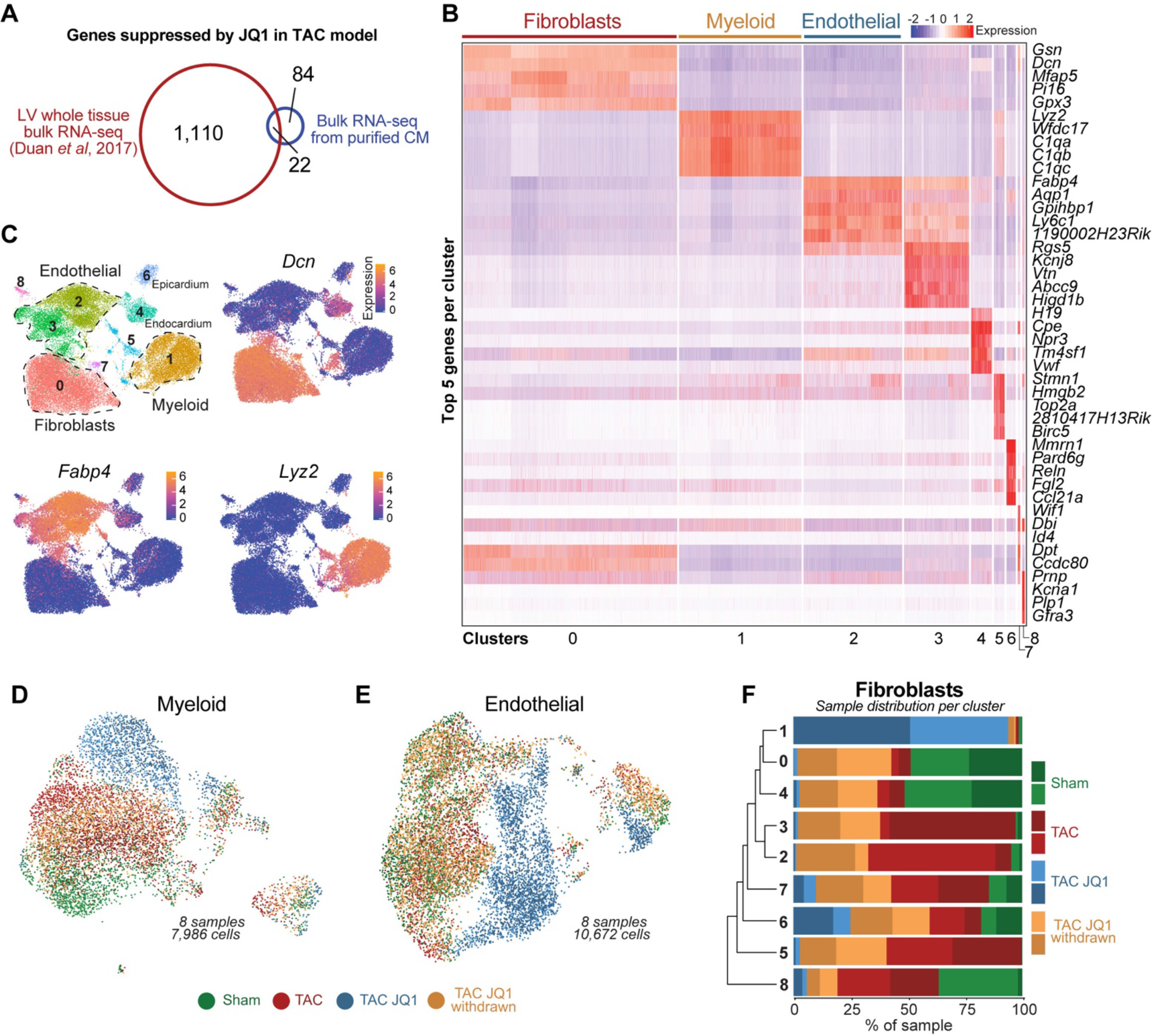
Single cell transcriptional landscape of non-cardiomyocytes in HF during intermittent exposure to BET bromodomain inhibition. **A.** Venn diagram showing overlap of TAC-induced and JQ1-suppressed genes (LogFC>0.5: pval<0.05) between bulk RNA-Seq from undissociated LV tissue (PMID 28515341) and extracted CMs. **B.** Heatmap showing the top 5 markers per cluster in the scRNAseq. Total cells n = 35,551 in 9 clusters. **C.** UMAP plots showing cluster identity of all cells (n = 35,551) and expression of 3 cell-population enriched markers (*Dcn* for FBs, *Lyz2* for Myeloid and *Fabp4* for endothelial) **D.** UMAP plot of myeloid cells subclustered colored by sample identity (n = 7,986). **E.** UMAP plot of endothelial cells subclustered colored by sample identity (n = 10,672). **F.** Histograms showing percentage of each sample in each FB cluster.

**Extended data Figure 2:**
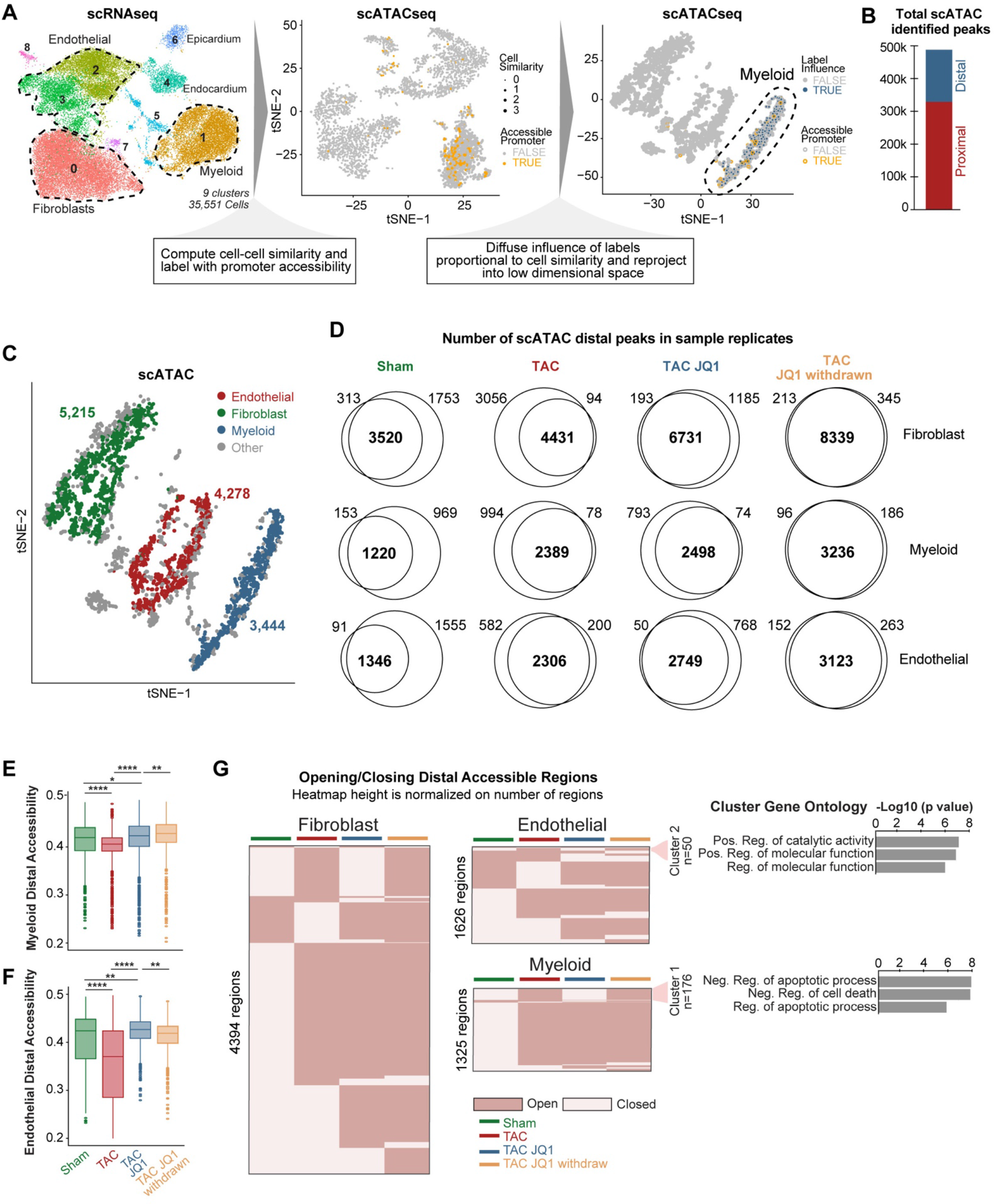
Single cell ATAC defines chromatin accessibility in HF during intermittent exposure to BET bromodomain inhibition. **A.** Schematic highlighting the approach to integrate scRNAseq with scATAC^21^. See extended methods for details. **B.** Total scATAC proximal and distal peaks identified in all cells. **C.** scATAC tSNE plot showing clusters and cell number of FBs, myeloid and endothelial cells after integration with scRNAseq. **D.** Venn diagrams showing sample replicate convergence of distal scATAC peaks. **E,F.** Chromatin accessibility at distal elements between samples in myeloid (e) and endothelial (f) cells. Trimming of 10% most extreme points was performed for better visualization. **G.** Dynamic accessibility of distal elements in FBs, myeloid and endothelial cells clustered by trend across samples. Top three GO terms for nearest genes to distal elements are shown for the cluster of accessible regions in myeloid and endothelial cells that are closed in Sham, opened in TAC, closed by JQ1, and reopened following JQ1 withdrawal.

**Extended data Figure 3:**
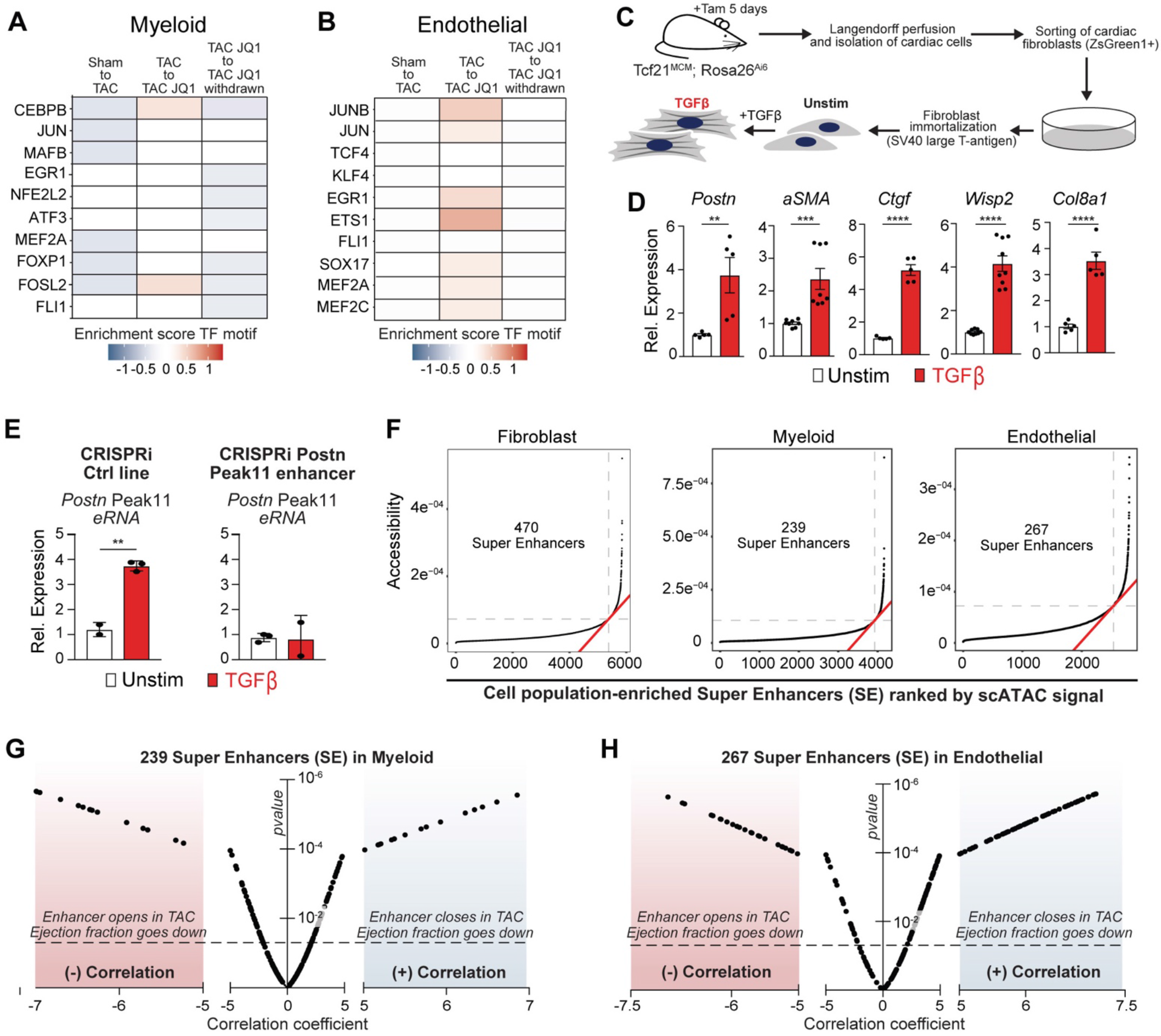
Characterization of a catalog of super enhancers in FBs, myeloid and endothelial cells. **A,B.** Enrichment scores for TF motif accessibility in distal elements between samples for the ten most expressed TFs in TAC in myeloid (a) and endothelial (b) cells. **C.** Schematic of isolation and immortalization of cardiac FBs used for subsequent PROseq, CRISPRi, 4C and ChIPseq experiments. **D.** Expression by qPCR of canonical markers of activated FBs in Unstim and TGFβ-treated cells. **E.** *Peak11 eRNA* expression measured by qPCR in Unstim and TGFβ-treated FBs in a CRISPRi control line versus CRISPRi targeting the Peak11 *Postn* enhancer (each panel is normalized to its Unstim condition). **F.** Distribution of accessibility in FB, myeloid and endothelial cells in TAC state identifies a class of distal regions (super enhancers, SE) where the accessibility falls over the inflection point of the curve. **G,H.** Volcano plots showing correlation coefficients and corresponding *p*-values (referred to analysis depicted in Fig. 2G) of 239 SEs in myeloid (g) and 267 SEs endothelial (h) cells. For D. and E., **P < 0.01, ***P < 0.001 and ****P < 0.0001 for indicated comparison. Data are shown as means ± SEM.

**Extended data Figure 4:**
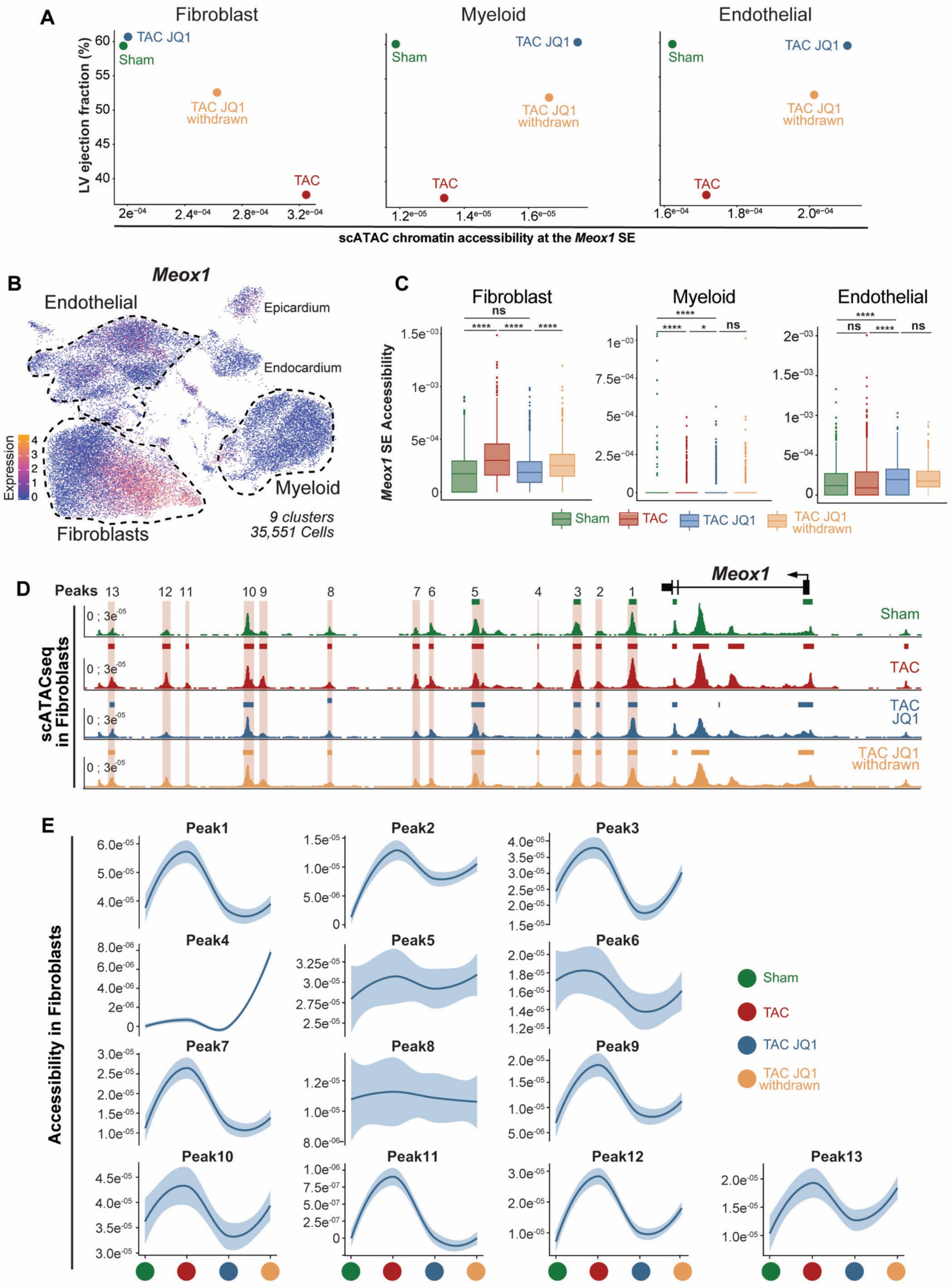
Chromatin accessibility at the Meox1 super enhancers is dynamic. **A.** Comparison of LV ejection fraction with chromatin accessibility at the *Meox1* SE in FBs, myeloid and endothelial cells. **B.** UMAP plot of *Meox1* expression in all cells (n=35,551). **C.** Chromatin accessibility at the *Meox1* SE between samples in FBs, myeloid and endothelial cells. **D.** scATAC coverage between samples at the *Meox1* SE in FBs identifies multiple dynamic peaks in HF with pulsatile BET inhibition. **E.** Chromatin accessibility trend between samples in all identified *Meox1* SE peaks. For C., *P < 0.05 and ****P < 0.0001 for indicated comparison. Data are shown as means ± SEM.

**Extended data Figure 5:**
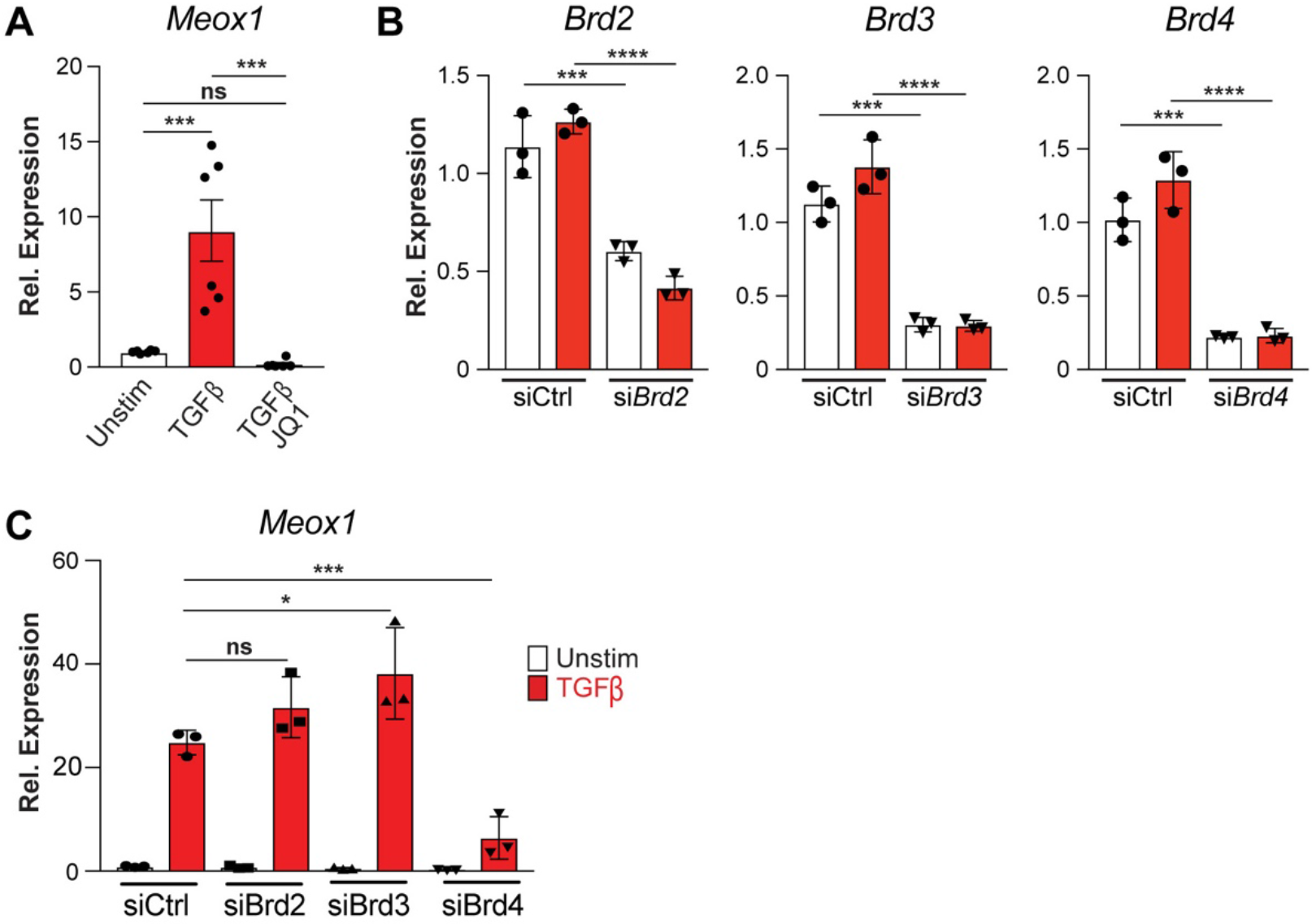
Brd4-dependent regulation of Meox1 expression. **A.** *Meox1* expression measured by qPCR in Unstim and TGFβ-treated FBs, with or without JQ1. **B.** Expression measured by qPCR of individual BET genes in Unstim or TGFβ-treated FBs with siRNA targeting either Ctrl, *Brd2, Brd3* or *Brd4.* **C.** *Meox1* expression measured by qPCR in Unstim or TGFβ-treated FBs with siRNA targeting either Ctrl, *Brd2, Brd3* or *Brd4.* For A-C., *P < 0.05, ***P < 0.001 and ****P < 0.0001 for indicated comparison. Data are shown as means ± SEM.

**Extended data Figure 6:**
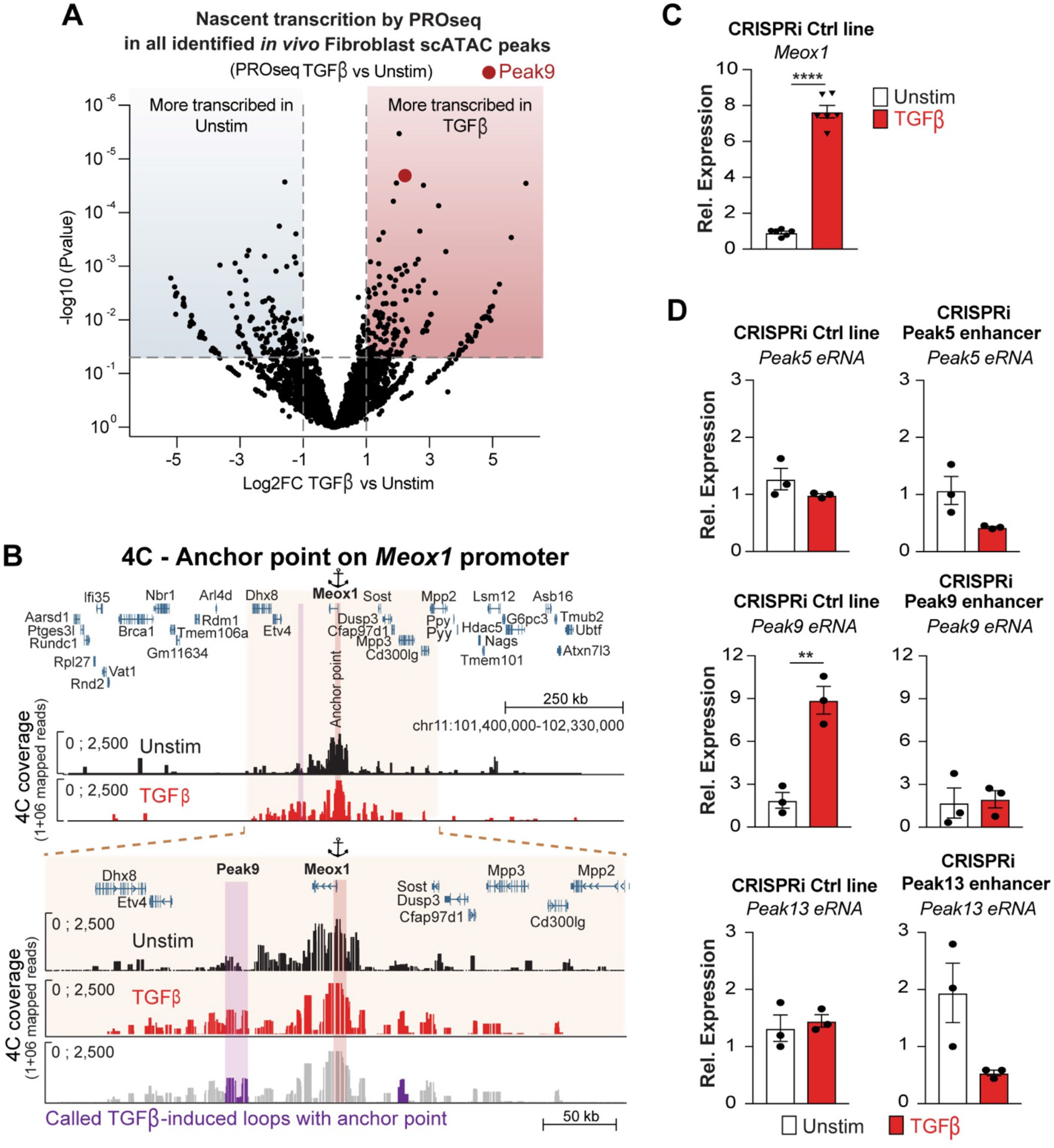
The Peak9 Meox1 enhancer is highly transcribed and contacts Meox1 promoter following TGFβ stimulation. **A.** Volcano plot showing LogFC of PROseq signal of all identified scATAC peaks in FBs (n=8163) between Unstim and TGFβ-treated FBs. **B.** Chromosome conformation capture (4C) between the *Meox1* promoter (anchor point) and Peak9 region 4C coverage in Unstim and TGFβ-treated FBs. A 922kb (top) genomic region and an expanded 328kb subset (bottom) are shown. The last track represents the called TGFβ-induced loops with the Peak9 (colored in purple). **C.** *Meox1* expression measured by qPCR in Unstim and TGFβ-treated in the CRISPRi control line. **D.** Expression measured by qPCR of Peak 5, 9 and 13 eRNAs in Unstim and TGFβ-treated conditions in the CRISPRi control line and the corresponding targeted CRISPRi lines (each panel is normalized to its Unstim condition). For C. and D. **P < 0.01 and ****P < 0.0001 for indicated comparison. Data are shown as means ± SEM.

**Extended data Figure 7:**
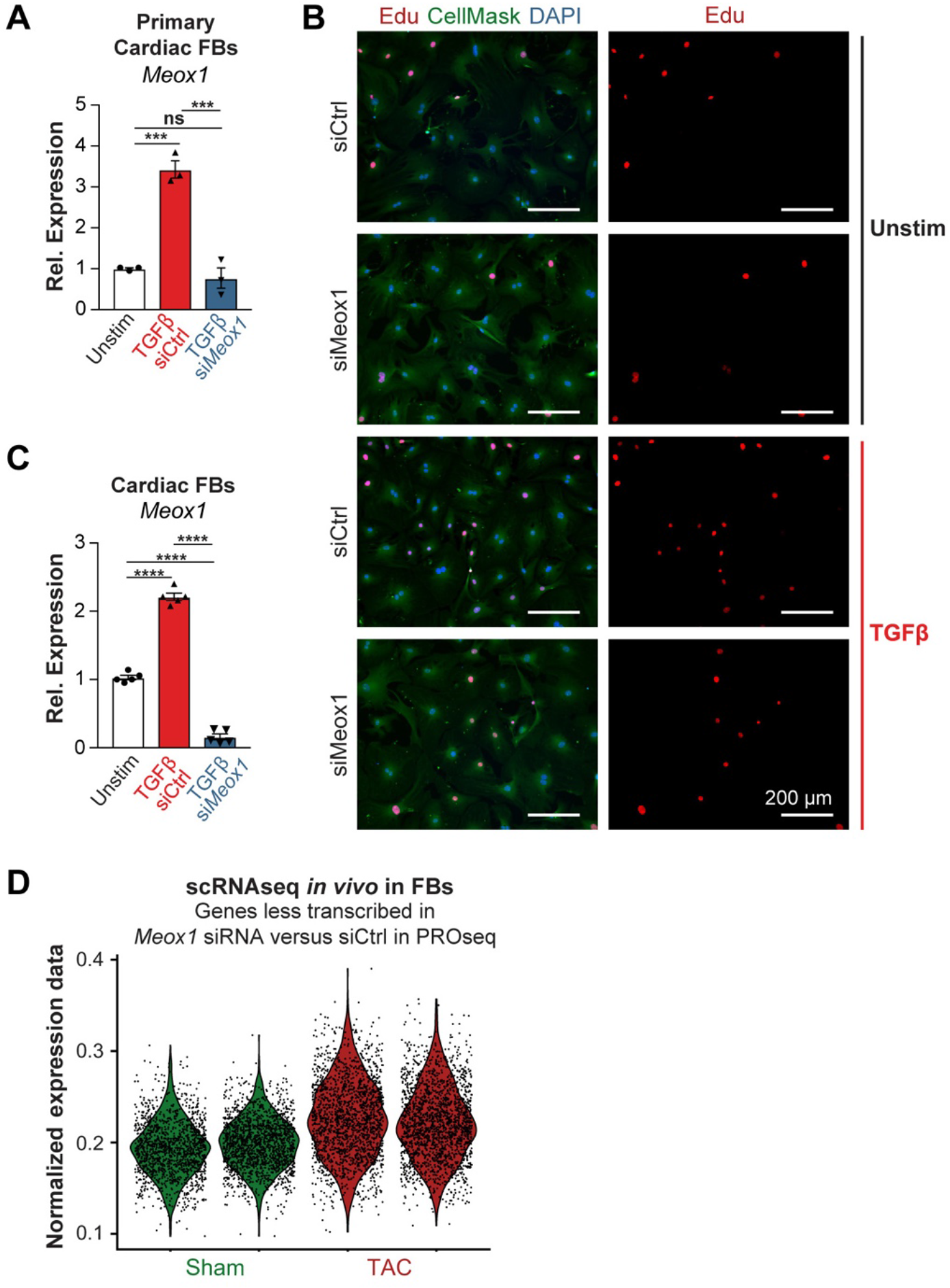
Meox1 controls the proliferative phenotype of FBs. **A.** *Meox1* expression measured by qPCR in mouse primary cardiac FBs in Unstim condition and TGFβ-treatment with siRNA targeting either Ctrl or *Meox1.* **B.** Representative images of Edu incorporation in Unstim (siCtrl and *siMeox1)* and TGFβ (siCtrl and *siMeox1)* conditions. DAPI (blue), Edu (green) and CellMask (red). **C.** *Meox1* expression measured by qPCR in immortalized cardiac FBs in Unstim condition and TGFβ-treatment with siRNA targeting either Ctrl or *Meox1.* **D.** Violin plot showing normalized expression score of a specific transcriptional signature (genes significantly less transcribed in *Meox1* siRNA versus siCtrl in PROseq that were captured in the scRNAseq *in vivo* - total number 340) in FB cells in Sham and TAC. For A. and C. ***P < 0.001 and ****P < 0.0001 for indicated comparison. Data are shown as means ± SEM.

## Methods

### Animal models

All protocols concerning animal use were approved by the Institutional Animal Care and Use Committees at the University of California San Francisco and conducted in strict accordance with the National Institutes of Health Guide for the Care and Use of Laboratory Animals. Studies were conducted with age-matched male C57Bl/6J (TAC/JQ1 related experiments) or CD1 mice (MI/CPI456 related experiments). Mice were housed in a temperature- and humidity-controlled pathogen-free facility with 12-hour light/dark cycle.

### CPI-456 dosing in mice post-myocardial infarction (MI)

CPI-456 was provided by Constellation Pharmaceuticals, Inc. A stock solution [175mg/ml CPI-456 in dimethyl sulfoxide (DMSO)] was diluted to working concentrations of 1mg/ml in 0.5% (w/v) methylcellulose. Homogeneous white suspension was achieved with sonication. CPI-456 was administrated by oral gavage with 10mg/kg following the schedule indicated in study. Vehicle control was an equal amount of DMSO dissolved in 0.5% (w/v) methylcellulose carrier solution. Dose formulation was prepared every four days, stored at 4°C when not in use.

### Mouse model of MI and echocardiography

MI was performed on adult CD1 mice of 7-8 weeks of age and between 25-30 grams of weight. The permanent proximal left anterior descending coronary artery ligation (LAD) was performed 2 mm distal to the inferior border of the left auricle using a 7-0 silk suture. Occlusion of arterial flow was demonstrated by ischemic color changes in the anterior wall of the heart. For sham surgeries, the LAD was surgically exposed without any further intervention. Echocardiography was performed as described^31^. The echocardiographer was blinded to the test groups. Mice were imaged using Vevo 3100 imaging system (FujiFilm VisualSonics Inc.) with RMV-707B 30-MHz probe. To minimize the confounding influence of different heart rates on left ventricular function, the flow of isoflurane (inhalational) was adjusted to anesthetize the mice while maintaining their heart rates at 450–550 beats per minute. LV ejection fraction was obtained from high-resolution two-dimensional measurements at the end-diastole and end-systole, standardized by twodimensional, long-axis views.

### Preparation of JQ1

JQ1 was synthesized and purified in the laboratory of Jun Qi (Dana-Farber Cancer Institute), as previously published^11^. For in vivo experiments, a stock solution [50 mg/ml JQ1 in dimethyl sulfoxide (DMSO)] was diluted to a working concentration of 5 mg/ml in an aqueous carrier (10% hydroxypropyl b-cyclodextrin; Sigma C0926) using vigorous vortexing. Mice were injected at a dose of 50 mg/kg given intraperitoneally once daily. Vehicle control was an equal amount of DMSO dissolved in 10% hydroxypropyl b-cyclodextrin carrier solution. All solutions were prepared and administered using sterile technique. For in vitro experiments, JQ1 was dissolved in DMSO and administered to cells at 500nM final concentration using an equal volume of DMSO as control.

### Mouse model of transverse aortic constriction and echocardiography

All mice were male C57Bl/6J mice aged 8-10 weeks from The Jackson Laboratory (Stock No: 000664). Mice were placed on a temperature-controlled small-animal surgical table to help maintain body temperature (37°C) during surgery. Mice were anesthetized with isoflurane, mechanically ventilated (Harvard Apparatus), and subjected to thoracotomy. For TAC surgery, the aortic arch was constricted between the left common carotid and the brachiocephalic arteries using a 7-0 silk suture and a 25-gauge needle, as previously described^3^. Intraperitoneal injections of JQ1 (50 mg/kg per day) or vehicle were administered 18 days after TAC surgery and continued as described in Figure 1c. For sham surgeries, thoracotomy was performed as above, and the aorta was surgically exposed without any further intervention. For echocardiography, mice were anesthetized with 1% inhalational isoflurane and imaged using the Vevo 3100 High Resolution Imaging System (FujiFilm VisualSonics Inc.) and the MX550S probe. Measurements were obtained from M-mode sampling and integrated electrocardiogram-gated kilohertz visualization (EKV) images taken in the LV short axis at the midpapillary level, as previously described^3^. LV areas and ejection fraction were obtained from high-resolution two-dimensional measurements at the end-diastole and end-systole, as previously described^3^.

### Langendorff perfusion and cells and nuclei isolation for subsequent single cell RNA and ATAC seq

Cell isolation from mouse hearts was performed as previously described with minor modifications^32^. Briefly, after proper anesthesia level was reached, thoracotomy was performed, and mouse heart was isolated. The isolated heart was cannulated and perfused with perfusion buffer (120.4 mM NaCl, 14.7 mM KCl, 0.6 mM KH2PO4, 0.6 mM Na2HPO4, 1.2 mM MgSO4, 10 mM Na-HEPES, 4.6 mM NaHCO3, 30 mM taurine, 10 mM 2,3-butanedione monoxime, and 5.5 mM glucose, pH 7.0) in a Langendorff perfusion system (Radnoti 120108EZ) for 5-10 minutes at 37 °C. The cannulated heart was then digested by digestion buffer (perfusion buffer with 300 units/mL collagenase II (Worthington Biochemical) and 50 μM CaCl2) for about 10 min at 37 °C. At the end of digestion, the atria and great vessels were removed and the ventricular tissue was transferred to and gently teased into small pieces in stop buffer (perfusion buffer with 10% fetal bovine serum) at 37 °C. After gently pipetting, cell suspension was passed through a 250uM strainer in a falcon tube and then at 30xg for 3 minutes at room temperature (RT). Then, the supernatant - containing most of the non-cardiomyocytes (CMs) - was divided from the pellet (containing the CM fraction). The non-CM fraction was centrifuged again at 30xg for 3 minutes at RT and the supernatant kept. The supernatant was then filtered with a cell strainer (70um) and finally centrifuged at 400xg for 3 minutes at RT for eliminating debris. The non-CM pellet was finally resuspended in 1mL cold PBS 0.5% BSA. 30k cells were counted with trypan blue using a hemocytometer and then used for subsequent 10X Genomics Chromium single cell RNAseq preparation. For single cell ATAC, 500k isolated and purified non-CMs were resuspended in 100uL lysis buffer (Tris-HCl 10mM pH 7.4, NaCl 10mM, MgCl2 3mM, Tween-20 0.1%, P40 0.1%, Digitonin 0.01%, BSA 1% in Nuclease-free water), pipetted 10 times and kept on ice for 5 minutes. Nuclei were then washed 1mL 1X PBS with 1% BSA and then centrifuged at for 5 minutes at 4C. The nuclei pellet was resuspended in 1mL 1X PBS with 1% BSA and filtered with a 10uM strainer (pluriSelect #43-50010-00). Nuclei were then counted with DAPI using a hemocytometer and finally 30k non-CM nuclei used for subsequent 10X Genomics Chromium single cell ATACseq preparation. The CM-fraction was, after the first centrifugation, centrifugated again at 30xg for 3 minutes at RT in stopping buffer and the supernatant was discarded. The CM pellet was finally centrifuged at 400xg for 3 minutes in stopping buffer at RT and after discarding the supernatant, the cell pellet was lysed in Qiazol (miRNeasy kit - Qiagen) for subsequent RNA extraction.

### Bulk RNA sequencing on purified cardiomyocytes

Total RNA from CMs was extracted using miRNeasy kit (Qiagen) according to the manufacturer’s instructions and quantified with Nanodrop (Thermo scientific). After RNA quality control with bioanalyzer Agilent 2100 (Agilent Technologies), Paired-end Poly(A)-enriched RNA libraries were prepared with the ovation RNA-seq Universal kit (NuGEN; strand specific) from the Gladstone Genomic core for 9 samples: Sham (x3), TAC-Veh (x3) and TAC-JQ1 (x3). High-throughput sequencing was done using a PE75 run on a NextSeq 500 instrument (Illumina). Reads were mapped to the mm10 reference mouse genome using STAR (v 2.7.3a) and assigned to Ensembl genes. We quantified gene expression using raw counts and kept the protein coding genes that showed an average FPKM value across the samples >0.5 FPKM. We then performed differential expression gene testing with DESeq2^33^ (v.1.24.0 R package) using default settings. Statistical significance was set at 5% false discovery rate (FDR; Benjamini-Hochberg).

### Single-cell transcriptome library preparation, sequencing and processing

Single-cell droplet libraries from the non-CM cell suspension (2x Sham, 2x TAC-Veh, 2x TAC-JQ1 and 2x TAC-JQ1 withdrawn) (Fig. 1c) were generated in the 10X Genomics Chromium controller according to the manufacturer’s instructions in the Chromium Single Cell 3’ Reagent Kit v.2 User Guide. Additional components used for library preparation include the Chromium Single Cell 3’ Library and Gel Bead Kit v.2 (PN-120237) and the Chromium Single Cell 3’ Chip kit v.2 (PN-120236). Libraries were prepared according to the manufacturer’s instructions using the Chromium Single Cell 3’ Library and Gel Bead Kit v.2 (PN-120237) and Chromium i7 Multiplex Kit (PN-120262). Final libraries were sequenced on the NextSeq 500 (Illumina) for a quality control run and then on the NovaSeq (Illumina) for deeper sequencing. All the 8 samples were pooled and sequenced in one single lane. Sequencing parameters were selected according to the Chromium Single Cell v.2 specifications. All libraries were sequenced to a mean read depth of at least 50,000 total aligned reads per cell. Raw sequencing reads were processed using the Cell Ranger v.2.2.0 pipeline from 10X Genomics. In brief, reads were demultiplexed, aligned to the mouse mm10 genome and UMI counts were quantified per gene per cell to generate a gene-barcode matrix. Data from the 8 samples were aggregated and normalized to the same sequencing depth, resulting in a combined gene-barcode matrix of all samples.

### Cell filtering and cell-type clustering for transcriptomic analysis

We sequenced the transcriptomes of 35,551 cells captured from our 8 samples (Fig. 1c). Filtering and clustering analyses of these cells were performed with the Seurat v.2.2 R package. Cells were normalized for genes expressed per cell and per total expression, then multiplied by a scale factor of 10,000 and log-transformed. Low quality cells were excluded from our analyses - this was achieved by filtering out cells with greater than 4,000 and fewer than 1,000 genes and cells with high percentage of mitochondrial genes (higher than 0.2%). Following the filtering step, we normalized the data (NormalizeData function, 10,000 default scale factor) and performed a linear regression on all genes (ScaleData function). We then performed a linear dimensional reduction (RunPCA function). Significant principal components were used for downstream graph-based, semi-unsupervised clustering into distinct populations (FindClusters function) and uniform manifold approximation and projection (UMAP)^34^ dimensionality reduction was used to project the cell population in two dimensions. For clustering, the resolution parameter was approximated based on the number of cells according to Seurat guidelines; a vector of resolution parameters was passed to the FindClusters function and the optimal resolution that established discernible clusters with distinct marker gene expression was selected. We obtained a total of 9 clusters representing the major adult cardiac non-CM cell populations. To identify marker genes driving each cluster, the clusters were compared pairwise for differential gene expression (FindAllMarkers function) using the Likelihood ratio test assuming an underlying negative binomial distribution (negbinom). We then isolated specific clusters (WhichCells function) for subsequent analysis on the fibroblast (cluster 0), myeloid (cluster 1) and endothelial (clusters 2 and 3) populations. Differential expression analysis between samples in the 3 major cell population was performed using the function diffExp=FindMarkers. For visualization of gene expression data between different samples a number of Seurat functions were used: FeaturePlot, VlnPlot and DotPlot. For the gene expression signature analysis in Extended data figure 7d, we took the 509 genes significantly less transcribed when *Meox1* was downregulated by siRNA versus siCtrl in the PROseq analysis, and then extract those genes that were also captured in our single cell RNAseq analysis (340/509). We then calculated a gene signature score by summing the mean expression of each of the 340 gene in every FB cells in Sham and TAC samples^35^. The resulting score was then plotted in Violin plot in Extended data figure 7d.

### Single-cell ATAC library preparation, sequencing and processing

After successful nuclei isolation, nuclei were then processed according to the 10X Genomics Single Cell ATAC kit v1.0 (PN-1000110) by first incubating with Tn5 Transposase for 1 hour, followed by GEM generation and barcode amplification using the 10X Genomics Chromium controller according to the manufacturer’s instructions. Additional components used for library preparation include the Chromium Chip E Single Cell ATAC kit (PN-1000082) and Chromium i7 Multiplex Kit N, Set A primers (PN-1000084). Final libraries were sequenced on the NextSeq 500 (Illumina) for a quality control run and then on the NovaSeq (Illumina) for deeper sequencing. All the 8 samples (2x Sham, 2x TAC-Veh, 2x TAC-JQ1 and 2x TAC-JQ1 withdrawn) (Fig. 1c) were pooled and sequenced in one NovaSeq single lane. Sequencing parameters were selected according to the Chromium Single Cell ATAC v1.0 specifications. All libraries were sequenced to a median read depth of at least 2,500 fragments/nuclei. Raw sequencing reads were processed using the Cell Ranger ATAC v1.0 pipeline from 10X Genomics. In brief, reads were demultiplexed and aligned to the mouse mm10 genome. As a test of sample quality, a minimum of 70% of fragments overlapped targeted regions as defined by CellRanger. Peaks are then are called on aggregated fragments and then barcodes with fewer fragments than an automatically determined threshold (usually around 200) within these peaks are discarded. The remaining fragments are counted to generate a peak-by-barcode matrix.

### Identifying cell types in single cell ATAC samples based on scRNA-seq

As previously described in the method, we determined a list of marker genes for each cluster in the scRNAseq data. Then, for each cell in each scATACseq sample, we computed the fraction of that cell’s accessible peaks that were in the promoters of each cluster’s marker genes. We computed a full cell-cell similarity matrix for each sample using Jaccard similarity of the binarized peak-by-cell matrix generated by CellRanger. Extended data figure 2a shows an example of mapping cluster 1 (Myeloid cells) from the scRNA-seq data to scATAC-seq sample. Cells marked in blue have accessible promoters for Myeloid marker genes. The size of each cell reflects how similar they are to a selected single cell. We then use the method described in Przytycki et al. to compute global influence scores of each label^21^. For each sample we chose a parameter *s* that maximized the median influence of labels corresponding to three cell types of interest (FB, Myeloid, and Endothelial) on cells that had accessible promoters for three selected marker genes *(Dcn, Lyz2, Fabp4* respectively). We assigned each cell a type based on the label with the highest influence on that cell. Extended data figure 2a shows that if we re-project cells into tSNE space using influence scores, cells with the same label cluster together (this projection is only used for illustrative purposes). Using this method, we identified 5,215 FB, 4,278 endothelial, and 3,444 as myeloid cells across eight samples.

### Determining cell type enriched peaks in single cell ATAC

To avoid bias, we computed cell type enriched peaks separately for each sample by comparing accessibility in cells of each type to cells of other types. For each cell type, to determine which peaks are cell type enriched, we repeatedly (ten times each) sampled a set of the same number of cells not of that type and with similar numbers of accessible peaks, and computed a one-tailed Wilcoxon test to determine if each peak was more accessible in the cell type being examined. We then combined sampled *p*-values using Fisher’s method and adjusted for multiple hypothesis correction using the Benjamini-Hochberg procedure. We considered a peak to be cell type enriched in a sample if it was significant at FDR < 0.1. For each cell type, we discarded any peaks that did not replicate in at least one other sample. This resulted in 22,467 FB enriched peaks, 12,602 endothelial enriched peaks, and 11,156 myeloid enriched peaks. We note that peaks can be enriched in more than once cell type.

### Computing single cell ATAC normalized accessibility scores

For each peak we computed a normalized accessibility score for each cell as one over the total number of accessible peaks in that cell. To compute overall accessibility trends in a cell type and condition, we calculated the fraction of all cell type enriched peaks in a cell that are accessible in that condition. For genome-wide normalized accessibility (e.g. for browser tracks) we instead normalized by number of fragments. To do this we first found all fragments that correspond to barcodes of cells that we want to calculate genome-wide accessibility for. We merge bed files for those cells using the unionBedGraphs function in bedtools2 to create a bedgraph. We assign an accessibility score to each region of the union bedgraph as the sum of number of fragments in each cell in that region over the total number of fragments for that cell over the number of cells used in the union.

### Calculating TF enrichment scores from single cell ATAC data

To assess the significance of changes in transcription factor binding between conditions for each cell type we trained a supervised learning model to link transcription factor binding locations to changes in gene expression. We used transcription factors (TF) with known vertebrate motifs included in HOMER, and selected the top 10 most expressed TFs in TAC in FB, myeloid and endothelial cells. First, for each transcription factor we determined all accessible binding sites in each condition using the “-find” option with the “findMotifsGenome.pl” command in HOMER using the set of distal cell type enriched peaks for that condition. We then generated an g-by-m matrix M^c^ for each condition c, where g is the number of genes and m is the number of transcription factors, by computing the distance from each binding site to the transcription start site of each gene. For each gene i and each binding location k for motif j, the corresponding entry in the matrix is defined as:

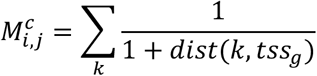

The difference in motif binding strength between two conditions c_1_ and c_2_ is then computed as *M*^*C*_1_^ – *M*^*C*_2_^. Then, given a vector Y of length g of log-fold change in expression of genes between the two conditions, we computed the importance of each transcription factor as the difference in change in expression for genes linked to that transcription factor minus genes not linked to that transcription factor while accounting for the effects of other transcription factors by using the targeted Maximum Likelihood Estimation (tMLE) approach described in Stone et al.^36^

### Generating atlas of super-enhancers and correlate chromatin accessibility with ejection fraction

To find super-enhancers, we first stitched together all cell type enriched peaks within 12.5kb of each other that were not separated by a gene using the single cell ATAC data from the TAC Veh samples. We then computed the normalized accessibility for each potential super-enhancer and used the ROSE algorithm^27^ to determine the threshold at which regions could be called superenhancers. We used this method to build a catalog of super-enhancers for FB, myeloid and endothelial cells in the diseased heart (TAC Veh). We then calculated how well each superenhancer’s accessibility correlates with left ventricle ejection fraction (EF). For each superenhancer we fit a linear model with a vector of EF observations across the four conditions as the response variable and the mean accessibility across the given cell type and across other cells as types as two term vectors. The fitted coefficient of the model for the given cell type is amount of change in EF explained by changes in accessibility in that cell type when controlling for changes in accessibility in other cell types. We call this value the correlation coefficient for each superenhancer for each cell type, with the significance of each coefficient determined by the p-value of that term in the linear model.

### Generation of cardiac FB immortalized cell line, culture condition and TGF-β stimulation

A Tcf21MCM^37^ mouse was crossed with a Rosa26-Ai6 mouse (Jackson Laboratory stock# 007906). At 10 weeks of age, intraperitoneal injection of Tamoxifen (75 mg tamoxifen/kg) was done for 5 days (once a day). Tamoxifen was prepared following Jackson Laboratory guidelines (https://www.jax.org/research-and-faculty/resources/cre-repository/tamoxifen#). After 5 days of injection, the mouse was sacrificed and the non-CM cells isolated as described in a previous section of this method. 100k ZsGreen positive cells were sorted with BD AriaII sorter and cultured for 3 days at 37°C in a humidified incubator with 5% CO2 and maintained in FB medium: high glucose DMEM (Life Technologies) supplemented with 10% fetal bovine serum (FBS) (Hyclone, GE Healthcare), 1x Non-Essential Amino Acid (NEAA), 10U/ml penicillin/streptomycin and 1mM sodium pyruvate (all from Life Technologies). For immortalization, FB were seeded at 10^5^ per 100 mm plate in the afternoon. Next morning, 2 hours before infection, fresh media was added on plates. The SV40 T Antigen Cell Immortalization Kit (Alstem, #CILV01) was used. 5 μl/well lentivirus supernatant (vials have 20ul) in the presence of Polybrene (added to a final concentration of 5 μg/ml). The next day, remove viral supernatant and switch again to FM medium. After 72 hours incubation, subculture the cells into 2 × 100 mm dishes and add the appropriate amount of puromycin for stable cell-line generation (1ug/ml for mouse cardiac FBs). After 10 days after puromycin selection, multiple clones were picked for expansion. For daily cell culture, FBs were split every 3-4 days and the media was changed every other day. Same media was used for HEK-293T cell culture for lentiviral production (see next sections). For the FB TGF-β stimulation, FBs were seeded at 1×10^5^/well of 6 well plate at day 1. On day 2, the media was changed to same basal media with 0.5% FBS. On the day 3, TGF-β 1(Peprotech) was added into the media at concentration of 10ng/ml. Cells were collected on day 5 for RNA extraction and subsequent gene expression analysis with real time PCR.

### RNA extraction, RT-PCR and real-time PCR analysis

Quantitative RT-PCR. Cells were harvested in TRIzolTM LS reagent (Invitogen) and total RNA was extracted using the Direct-Zol RNA kit (Zymo Research) according to manufacturer instruction. 500ng of RNA was converted to cDNA using SuperScriptTM III First-strand Synthesis SuperMix for qRT-PCR (Invitrogen). For Taqman real-time PCR, 1/50 cDNA was applied for quantitative PCR reaction using Taqman Universal PCR master mix (Life technologies). The PCR was conducted in 7900HT Fast Real-Time system (Applied Biosystem). The Taqman probes are listed in the ‘Taqman probes table’. For eRNA expression analysis, 1/30 cDNA was applied for quantitative PCR reaction using SsoAdvanced Universal SYBR green supermix (Bio-Rad). Primer sequences are listed in the ‘Syber primers for enhancer RNAs (eRNAs) table’. All gene expressions were normalized with *Actb* gene.

### Precision nuclear run-on sequencing (PROseq)

PRO-seq experiments were performed as previously reported^38^ with a few modifications. Briefly, 3 million Cardiac FBs were cultured as described previously in this method. After 48h of TGF-β treatment, cells were washed 3 times with cold PBS and then sequentially swelled in swelling buffer (10mM Tris-HCl pH7.5, 2mM MgCl2, 3mM CaCl2) for 10 min on ice, harvested, and lysed in lysis buffer (swelling buffer plus 0.5% NP-40, 20 units of SUPERase-In, and 10% glycerol). The resultant nuclei were washed two more times with 5ml lysis buffer and finally resuspended in 100 μl of freezing buffer (50mM Tris-HCl pH8.3, 40% glycerol, 5mM MgCl2, 0.1mM EDTA). For the run-on assay, resuspended nuclei were mixed with an equal volume of reaction buffer (10 mM Tris-HCl pH 8.0, 5 mM MgCl2, 1 mM DTT, 300 mM KCl, 20 units of SUPERase-In, 1% sarkosyl, 100 M A/GTP, 100 μM biotin-11-C/UTP (Perkin-Elmer) and incubated for 5 min at 30°C. The resultant nuclear-runon RNA (NRO-RNA) was then extracted with TRIzol® LS reagent (Life Technologies, Cat# 10296–028) following manufacturer’s instructions. NRO-RNA was fragmented to 200–500nt by alkaline base hydrolysis on ice for 30 min and neutralized by adding 1× volume of 1 M Tris-HCl pH 6.8, Excessive salt and residual NTPs were removed by using P30 column (Bio-Rad, Cat# 732–6250), followed by treatment with DNase I (Promega Cat# M6101) and antarctic phosphatase (NEB Cat# M0289L). Fragmented nascent RNA was bound to 10 μl of MyOne Streptavidin C1 dynabeads (Invitrogen, Cat# 65001) following the manufacturer’s instructions. The beads were washed twice in high salt (2 M NaCl, 50 mM Tris-HCl pH 7.5, 0.5% Triton X-100, 0.5 mM EDTA), once in medium salt (1M NaCl, 5 mM Tris-HCl pH 7.5, 0.1% Triton X-100, 0.5 mM EDTA), and once in low salt (5 mM Tris-HCl pH 7.5, 0.1% Triton X-100). Bound RNA was extracted from the bead using Trizol (Invitrogen, Cat# 15596–018) in two consecutive extractions, and the RNA fractions were pooled, followed by ethanol precipitation. Libraries were generated using the NEBNext® Multiplex Small RNA Library Prep Set. (NEB, Cat#E7300S) following the manufacturer’s instructions. The cDNA products were separated on a 10% polyacrylamide TBE-urea gel and only those fragments migrating between 200–500bp were excised and recovered by gel extraction. Finally, libraries were quantified by Qubit and sent to sequence SR75 bp on a HiSeq 4000 platform (Illumina). Results coming from the deep sequencing have been analyzed as follow: low quality reads have been removed using Trimmomatic, remaining reads have been aligned to mm10 reference genome using bowtie2, differential expression and any downstream analysis have been performed using homer.

### CRISPR interference (CRISPRi) for sequence-specific repression

For repressing enhancer activity, CRISPRi was used. The lentiviral plasmid, pHR-SFFV-KRAB-dCas9-mcherry (gift from Dr. Jonathan Weissmen, Addgene: 60954) was used. For constructing gRNA lentiviral vector, we modified pU6-sgRNAEF1Alpha-puroT2A-BFP (gift from Dr. Jonathan Weissmen, Addgene: 60955) by replacing the puromycin gene with Hygromycin gene and made pU6-sgRNAEF1Alpha-HygT2A-BFP. Pairs of synthesized gRNA oligos (‘CRISPRi guide RNAs targeting enhancers table’) with 5’ and 3’ overhangs were annealed and sub-cloned into BstXI and BlpI double digested pU6-sgRNAEF1Alpha-HygT2A-BFP by T4 ligase mediated ligation. The construct was sequencing verified (Quintara Bio, Berkeley, CA, USA). The gRNAs for repressing enhancer peaks were chosen by the program Chopchop (http://chopchop.cbu.uib.no). For generating the lentiviral particles, 2×10^6^ HEK-293T cells were seeded on a 100mm plate one day prior the transfection and cultured in 10ml FB media. On the day of transfection, the old media was replenished with 8ml of fresh media, then 5μg of desired lentiviral vector was co-transfected with 2.5μg of envelope protein vector pMD2.G (Addgene:12258), and 2.5μg of the packaging vector psPAX2 (Addgene: 12260) into HEK 293T cells using 59ul of FUGENE HD transfection reagent (Promega, San Luis Obispo, CA, USA) following the manufacturer’s instruction. 48 hours after transfection, supernatant was collected and 12ml of fresh media was added to the plate and culture for another 24 hours for secondary collection of the supernatant. All supernatants were filtered through 45μm PVDF syringe filter (Thermo Fisher Scientific) to remove the cell debris contamination. The obtained supernatant was then used for performing transduction on cardiac FBs. Firstly, we generated a FB CRISPRi control line. We transduced our FBs with lentiviral vector that expressed the KRAB-dCas9-mcherry. We then collected the FBs and single cell sorted in to 96 well plate by flow cytometry (BD Arial II) and generated a clonal FB line expressing the CRISPRi machinery. The expression of the dCas9 was confirmed by western blot (data not shown) and the FB line with the highest dCas9 expression (referred as clone C1) was used for subsequent experiments. For generating the FB lines with targeted repression of the enhancer regions in this study, the C1 clone was transduced with three gRNA viral vectors that target the desired region (one gRNA targeting the center of the enhancer peak, the other targeting the two extreme parts of the peak). Pure polyclonal population of targeting gRNAs were selected by hygromycin selection at 200μg/ml for 7 days. These gRNA-C1 cell lines were then used together with the C1 control line for TGF-β stimulation (see other section of this method).

### Generation of MEOX1-HA cardiac FB line

The pHR-HAtag-mMeox1 vector was constructed by PCR amplifying the HA tag-mMeox1 fragment from the vector HA-tag-MEOX1 mouse (Twist Biosciences). For generating the pHR-HAtag-mMEOX11, we replaced the KRAB and dCas9 cassettes from the pHR-SFFV-KRAB-dCas9-mCherry vector with the HA-tag-MEOX1 cassette using Cold-fusion cloning kit (SBI System BioSciences, Palo Alto, CA, USA) following the instruction provided by the manufacturer. The construct was sequencing verified. For generating the lentiviral particles, the same procedure described in the section “CRISPR interference (CRISPRi) for sequence-specific repression” was used. The obtained supernatant was then used for performing transduction on cardiac FBs with the lentiviral vector that express HA-tag-MEOX1 and mCherry. Pure polyclonal population of HA-tag-MEOX1 FBs were sorted by flow cytometry (BD Arial II) for stable mCherry expression. This HA-tag-MEOX1 FB line was used for subsequent chromatin immunoprecipitation followed by sequencing (ChIPseq).

### ChIPseq assay, library preparation and analysis

For ChIPseq experiment, 10×10^6^ HA-tag-MEOX1 FBs were pelleted and suspended in 10ml DMEM and cross-linked in 1% formalin solution (Thermo Fisher Scientific) by rocking in room temperature for 10 minutes. Then glycine (final concentration 0.125M) was added to quench the cross-link for 5minutes. Samples were centrifuged at 1000 rcf for 5minutes at 4°C. Cells were washed with 10ml of cold 1x PBS supplemented with proteinase inhibitors and phosphatase inhibitors (Roche # 4693132001) and the pellets were snap frozen in liquid nitrogen. All samples were stored at −80°C until use. When ready, cell pellets were incubated in cell lysis buffer (20 mM Tris-HCl, pH 8, 85 mM KCl, 0.5% NP-40, protease inhibitors for 10 min on a rotator at 4°°C. Nuclei were isolated by centrifugation (2,500 x g, 5 min, 4°C), resuspended in nuclear lysis buffer (50 mM Tris-HCl, pH 8, 10 mM EDTA, pH 8, 1% SDS, protease inhibitors) and incubated on a rotator for 30 min at 4°C. Chromatin was sheared using a Covaris S2 sonicator (Covaris Inc) for 20 min (60 s cycles, 20% duty cycle, 200 cycles/burst, intensity = 5) until DNA was in the 200–700 basepair range. Chromatin was diluted 3-fold in ChIP dilution buffer (0.01% SDS, 1.1% Triton X-100, 1.2mMEDTA, 16.7mMTris-HCl, pH 8, 167 mM NaCl, protease inhibitors) and incubated with 3ul of anti-HA antibody (Abcam #9110) at 4°C overnight under rotation. Antibody-protein complexes were immunoprecipitated using Pierce Protein A/G magnetic beads at 4°C for 2 h under rotation. Beads were washed five times (2-min/wash under rotation) with cold RIPA buffer (50 mM HEPES-KOH, pH 7.5, 500 mM LiCl, 1 mM EDTA, 1% NP-40, 0.7% Na-deoxycholate), followed by one wash in cold final wash buffer (1xTE, 50 mM NaCl). Immunoprecipitated chromatin was eluted at 65°C with agitation for 30 min in elution buffer (50mMTris-HCl pH 8.0, 10mMEDTA, 1% SDS). High-salt buffer (250mM Tris-HCl, pH 7.5, 32.5 mM EDTA, pH 8, 1.25M NaCl) and Proteinase K (New England Biolabs Inc (NEB)) were added and crosslinks were reversed overnight at 65°C. Samples were treated with RNase A, and DNA was purified with AMPure XP beads (Beckman Coulter cat #A63881). Fragmented ChIP and input DNA were end-repaired, 5’-phosphorylated and dA-tailed with NEBNext Ultra II DNA Library Prep Kit for Illumina (NEB, E7645). Samples were ligated to adaptor oligos for multiplex sequencing (NEB, E7335), PCR amplified, and sequenced on an Illumina NextSeq 500 at the Gladstone Institutes.

For the ChIPseq analysis, trimming of known adapters and low-quality regions of reads was performed using Fastq-mcf. Sequence quality control was assessed using FastQC (http://www.bioinformatics.babraham.ac.uk/projects/fastqc/). Alignment to the mm10 reference genome was performed using Bowtie 2.2.4^39^. Peaks were called using GEM^40^. Read counts per peak were generated with featureCounts^41^ and normalized to account for differences in sequencing depth between samples using upper quartile normalization separately for the ChIP and input sample. Regions bound by MEOX1 were determined using empirical Bayes F-tests for a quasi-likelihood negative binomial generalized log-linear model of the count data as implemented in edgeR. Specifically, we tested for a significant (i.e., non-zero at FDR < 5%) log2 fold-increase in normalized peak signal for ChIP versus the corresponding input sample. 3 separate samples (and relative inputs) were ran from 3 different ChIP assays and only peaks present in minimum 2 out of 3 replicates were kept. Specifically, region intersections were found using BEDTools^42^.

### Circularized Chromosome Conformation Capture (4C)

4C-seq experiments were largely performed as previously reported^43^ with a few modifications. Briefly, 10 million Cardiac FBs were cultured as previously described in this method. After 48h of TGF-β treatment, cells were washed 3 times with cold PBS and then sequentially cross-linked with 1% formaldehyde for 10 min. The reaction was quenched by addition of glycine to a final concentration of 0.125 M. Cells were washed once again with cold PBS and nuclei were extracted by incubation 10min in ice with 5 ml of ice-cold lysis buffer (10 mM Tris–HCl, pH 8.0, 10 mM NaCl, 0.2% NP-40 + protease inhibitor). Nuclei were resuspended in restriction enzyme buffer (NEB Cat#B7006S) and incubated with 0.3 % SDS for 1h at 37 °C and further incubated with 2% Triton X-100 for 1h. 400U of DpnII restriction enzyme (NEB Cat#R0543S) was added and incubated overnight. Restriction enzyme was heat inactivated at 65°C for 20 min. Ligation of DNA regions in close physical proximity was performed using 1000U of T4 DNA ligase (NEB M0202M) overnight. The following day, 300μg of proteinase K were added and decrosslinking was performed at 65°C overnight. After de-crosslinking, DNA was purified using phenol/chloroform precipitation and the second digestion was performed by adding 400U of NlaIII restriction enzyme (NEB Cat#R0125S), then incubation at 65°C for 4h. After the second digestion, ligation of DNA was performed again using T4 DNA ligase as described above. 4C-seq libraries were amplified using PCR with primer containing partial Illumina sequence adaptors (1^st^ primer: gttcagagttctacagtccgacgatc; 2^nd^ primer: agacgtgtgctcttccgatct). The first primer was designed on each view point and the second primer designed beside the closest NlaIII cutting site to the view point. The primer sequences used are listed in the ‘4C primers table’.

Full-length Illumina sequencing adaptors and barcodes were added by the second round of PCR. Finally, libraries were quantified by Qubit and library quality checked with bioanalyzer Agilent 2100 (Agilent Technologies). High-throughput sequencing was done using a SE75 run on a NextSeq 500 instrument (Illumina). Sequencing results have been analyzed as previously described^44^. Peak calling over background has been performed using de function “doPeakC” directly on the rds files produced by the pipeline as described^44^.

### siRNA transfection on cardiac FBs

1×10^5^ of immortalized FBs or primary adult cardiac FBs were seeded before the day of transfection in 6 well plate. On the day of transfection, the wells for the subsequent TGB-β treatment were switched on FB medium with 0.5% FBS, while the Unstimulated wells were switched with normal FB medium with 10% FBS. Cells were transfected with either 15nM mouse Meox1 siRNA (Thermo fisher) or 15nM siRNA Control, using 7μl of Lipofectamine™ RNAiMAX transfection reagent (Thermo Fisher) for each well according to the manufacturer’s instruction. 24 hours after transfection, the wells with 0.5% FBS medium were treated with TGF-β1 (detailed procedure described in another section of this method).

### Primary adult cardiac fibroblast isolation and culture

Primary adult mouse ventricular fibroblasts (AMVFs) were prepared with minor modifications to the protocol described previously^45^. Briefly, mice were anesthetized with isoflurane and administered 100 μL of heparin (100 U/mL) via intraperitoneal injection. The hearts were excised and immediately suspended on a Langendorff apparatus by cannulation of the aortic root and perfused at a constant rate of 4 mL/min at 37C starting with 4 minutes of perfusion buffer (113 mM NaCl, 4.7 mM KCl, 0.6 mM KH2PO4, 0.6 mM Na2HPO4, 1.2 mM MgSO4, 10 mM HEPES, 12 mM NaHCO3, 10 mM KHCO3, 30 mM Taurine, 10 mM 2,3-Butanedione monoxime, 5.5 mM D-(+)-glucose, pH 7.4). Subsequently, enzymatic digestion was achieved by 3 min of perfusion with calcium-free digestion buffer (400 units/mL of collagenase II in perfusion buffer; Worthington LS004177) followed by 12 min of perfusion with digestion buffer containing 50 μM CaCl2. Hearts were removed from the perfusion apparatus, atria were removed and ventricles placed in Stopping Buffer (10% FBS and 12.5 μM CaCl2 in perfusion buffer). Ventricles were gently mechanically disrupted using transfer pipettes until tissue was sufficiently digested. The cell suspension was filtered through a 250 μm mesh and CMs were allowed to settle by gravity for 10 min; the supernatant, containing the first non-CM fraction, was collected. CMs were resuspended in an additional 10 mL Stopping Buffer and subsequently allowed to settle for 10 minutes. Supernatant was collected and both non-CM fractions were centrifuged at 500 xg for 5 min. CMs were discarded. Non-CMs were resuspended, combined and plated in growth medium consisting of DMEM/F12 media (Corning 10-092-CV) supplemented with 10% BenchMark™ FBS (Gemini Bio-Products 100-106), 1% Penicillin Streptomycin L-Glutamine (Corning 30-009-Cl) and 1 μmol/L ascorbic acid. Upon reaching 80% confluency, AMVFs were passaged once to P1, in an attempt to deter spontaneous activation, and plated appropriately for downstream assays.

### Collagen gel contraction assay

Compressible collagen matrices were prepared in 24-well plates using PureCol EZ Gel Solution (Advanced BioMatrix 5074) by incubating at 37C for 1.5 hrs. Passage 1 AMVFs suspended in serum-supplemented growth medium were seeded (150,000 cells/gel) on the collagen gels for 24 hrs prior to equilibration by serum deprivation (0.1% FBS) overnight. During serum-starvation, cells were also transfected with an siRNA directed against murine Meox1 (or a negative control siRNA) using Lipofectamine™ RNAiMAX Transfection Reagent (ThermoFisher Scientific 13778030) according to the manufacturer’s instructions. At the initiation of contraction, gels were released from the walls of the well, transfection reagent was removed and cells were treated with 10 ng/mL TGF-β1 (Fisher Scientific 50725143) for 72 hrs. Well images were captured every 24 hrs; gel area for each well was determined using ImageJ software and data are reported as percent contraction.

### EdU Incorporation Assay

Passage 1 AMVFs were seeded in 12-well plates containing glass coverslips at a density of 20,000 cells/well for 24 hrs, followed by equilibration in serum-starvation media (0.1% FBS) overnight. During serum-starvation, cells were also transfected with an siRNA directed against murine Meox1 (or a negative control siRNA) using Lipofectamine™ RNAiMAX Transfection Reagent (ThermoFisher Scientific 13778030) according to the manufacturer’s instructions. Media was exchanged and cells were stimulated with 10 ng/mL TGF-β1 (Fisher Scientific 50725143) for 48 hrs; AMVFs were simultaneously incubated with 10 μM 5-ethynyl-2’-deoxyuridine (EdU) to label proliferative cells over the 48 hr period of stimulation. AMVFs were fixed in 3.7% formaldehyde and EdU-positive cells were detected using the Click-iT® EdU Imaging Kit for Imaging (ThermoFisher Scientific C10337) according to the manufacturer’s protocol. Coverslips were mounted using ProLong™ Diamond Antifade Mountant (Invitrogen P36961) and allowed to cure overnight prior to imaging on a Keyence BZ-X710 fluorescence microscope. Percent EdU incorporation was determined by quantifying the percent of EdU positive cells relative to the number of nuclei detected per field.

### Gene Ontology (GO) analysis

GO analysis on distal elements was performed using GREAT (http://great.stanford.edu/public/html/). GO analysis on differential expressed protein coding genes was performed using Enrichr^46^.

### Statistics and reproducibility

Standard statistical analyses were performed using GraphPad Prism 8. When several conditions were to compare, we performed a one-way ANOVA, followed by Tukey range test to assess the significance among pairs of conditions. When only two conditions were to test, we performed Student’s t-test. All the p-values related to the figures showing chromatin accessibility were obtained with a two-tailed wilcoxon tests. For all quantifications related to cardiac function, gene expression by RT-qPCR, collagen contraction and EdU incorporation, the means ± SEM are reported in the figures. For gene expression data by RT-qPCR analysis or RNAseq FPKM, the number of replicates is indicated as data points in the graphs. The level of significance in all graphs is represented as follow: * P<0.05, ** P<0.01, *** P<0.001, **** P<0.0001.

Taqman probes table

**Taqman probes table**
| Gene | ThermoFisher Ref. # |
| --- | --- |
| *Actb* | Mm01205647 |
| *Meox1* | Mm00440285 |
| *Postn* | Mm01284919 |
| *Col8a1* | Mm01344185 |
| *Ctgf* | Mm01192933 |
| *Wisp2* | Mm00497471 |
| *Brd2* | Mm01271171 |
| *Brd3* | Mm01333576 |
| *Brd4* | Mm00480394 |

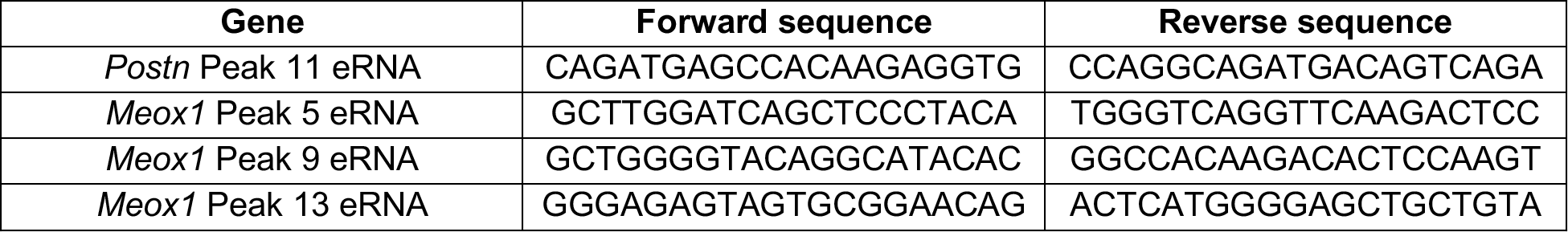
Syber primers for enhancer RNAs (eRNAs) table

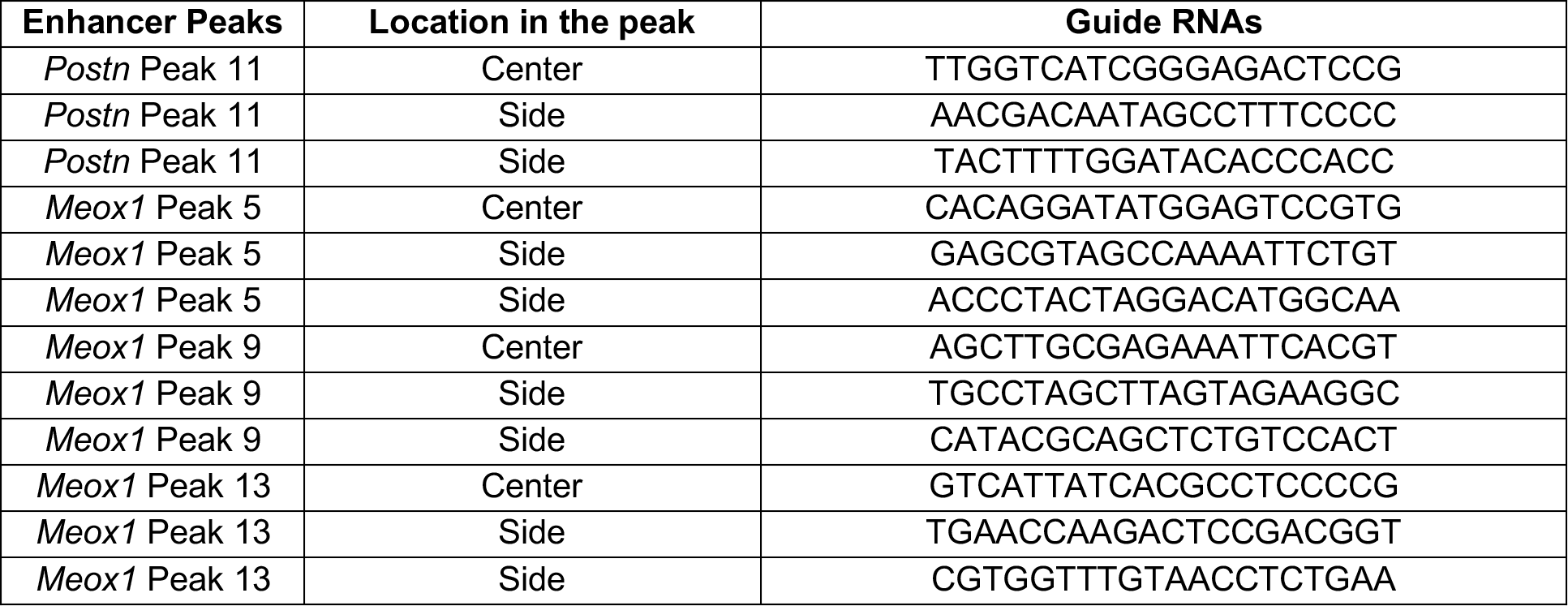
CRISPRi guide RNAs targeting enhancers table

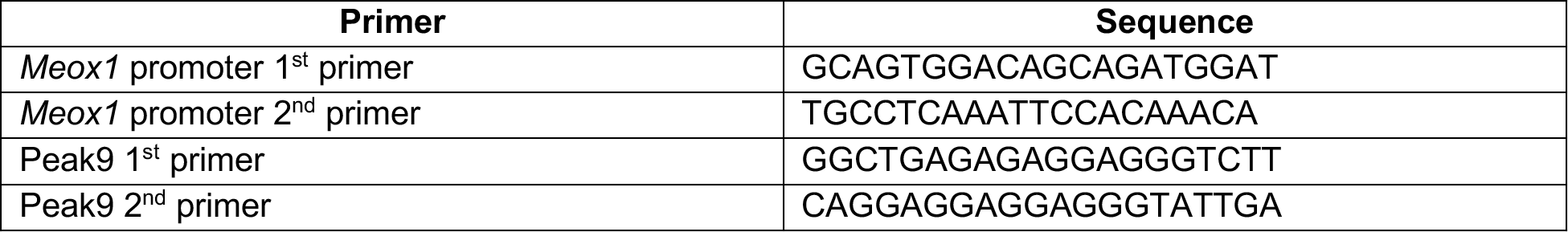
4C primers table

## Bibliography

1. Rockey, D. C., Bell, P. D. & Hill, J. A. Fibrosis--A Common Pathway to Organ Injury and Failure. N. Engl. J. Med. 373, 96 (2015).

2. Jun, J.-I. & Lau, L. F. Resolution of organ fibrosis. J. Clin. Invest. 128, 97–107 (2018).

3. Anand, P. et al. BET bromodomains mediate transcriptional pause release in heart failure. Cell 154, 569–582 (2013).

4. Spiltoir, J. I. et al. BET acetyl-lysine binding proteins control pathological cardiac hypertrophy. J. Mol. Cell Cardiol. 63, 175–179 (2013).

5. Duan, Q. et al. BET bromodomain inhibition suppresses innate inflammatory and profibrotic transcriptional networks in heart failure. Sci. Transl. Med. 9, (2017).

6. Stratton, M. S. et al. Dynamic chromatin targeting of BRD4 stimulates cardiac fibroblast activation. Circ. Res. 125, 662–677 (2019).

7. Antolic, A. et al. BET bromodomain proteins regulate transcriptional reprogramming in genetic dilated cardiomyopathy. BioRxiv (2020). doi:10.1101/2020.02.09.940882

8. Stratton, M. S. et al. Signal-Dependent Recruitment of BRD4 to Cardiomyocyte Super-Enhancers Is Suppressed by a MicroRNA. Cell Rep. 16, 1366–1378 (2016).

9. Virani, S. S. et al. Heart Disease and Stroke Statistics-2020 Update: A Report From the American Heart Association. Circulation 141, e139–e596 (2020).

10. Shi, J. & Vakoc, C. R. The mechanisms behind the therapeutic activity of BET bromodomain inhibition. Mol. Cell 54, 728–736 (2014).

11. Filippakopoulos, P. et al. Selective inhibition of BET bromodomains. Nature 468, 1067–1073 (2010).

12. Nicodeme, E. et al. Suppression of inflammation by a synthetic histone mimic. Nature 468, 1119–1123 (2010).

13. Taniguchi, Y. The Bromodomain and Extra-Terminal Domain (BET) Family: Functional Anatomy of BET Paralogous Proteins. Int. J. Mol. Sci. 17, (2016).

14. Albrecht, B. K. et al. Identification of a Benzoisoxazoloazepine Inhibitor (CPI-0610) of the Bromodomain and Extra-Terminal (BET) Family as a Candidate for Human Clinical Trials. J. Med. Chem. 59, 1330–1339 (2016).

15. Lovén, J. et al. Selective inhibition of tumor oncogenes by disruption of super-enhancers. Cell 153, 320–334 (2013).

16. Brown, J. D. et al. BET bromodomain proteins regulate enhancer function during adipogenesis. Proc. Natl. Acad. Sci. USA 115, 2144–2149 (2018).

17. Prabhu, S. D. & Frangogiannis, N. G. The biological basis for cardiac repair after myocardial infarction: from inflammation to fibrosis. Circ. Res. 119, 91–112 (2016).

18. Fu, X. et al. Specialized fibroblast differentiated states underlie scar formation in the infarcted mouse heart. J. Clin. Invest. 128, 2127–2143 (2018).

19. Kanisicak, O. et al. Genetic lineage tracing defines myofibroblast origin and function in the injured heart. Nat. Commun. 7, 12260 (2016).

20. Buenrostro, J. D., Wu, B., Chang, H. Y. & Greenleaf, W. J. ATAC-seq: A Method for Assaying Chromatin Accessibility Genome-Wide. Curr. Protoc. Mol. Biol. 109, 21.29.1–21.29.9 (2015).

21. Przytycki, P. F. & Pollard, K. S. Semi-supervised identification of cell populations in singlecell ATAC-seq. BioRxiv (2019). doi:10.1101/847657

22. Lu, D. et al. Meox1 accelerates myocardial hypertrophic decompensation through Gata4. Cardiovasc. Res. 114, 300–311 (2018).

23. Li, W., Notani, D. & Rosenfeld, M. G. Enhancers as non-coding RNA transcription units: recent insights and future perspectives. Nat. Rev. Genet. 17, 207–223 (2016).

24. Mahat, D. B. et al. Base-pair-resolution genome-wide mapping of active RNA polymerases using precision nuclear run-on (PRO-seq). Nat. Protoc. 11, 1455–1476 (2016).

25. Dobaczewski, M., Chen, W. & Frangogiannis, N. G. Transforming growth factor (TGF)-β signaling in cardiac remodeling. J. Mol. Cell Cardiol. 51, 600–606 (2011).

26. Gilbert, L. A. et al. CRISPR-mediated modular RNA-guided regulation of transcription in eukaryotes. Cell 154, 442–451 (2013).

27. Whyte, W. A. et al. Master transcription factors and mediator establish super-enhancers at key cell identity genes. Cell 153, 307–319 (2013).

28. Parker, S. C. J. et al. Chromatin stretch enhancer states drive cell-specific gene regulation and harbor human disease risk variants. Proc. Natl. Acad. Sci. USA 110, 17921–17926 (2013).

29. Litvinukova, M. et al. Cells and gene expression programs in the adult human heart. BioRxiv (2020). doi:10.1101/2020.04.03.024075

30. Sivakumar, P. et al. RNA sequencing of transplant-stage idiopathic pulmonary fibrosis lung reveals unique pathway regulation. ERJ Open Research 5, (2019).

31. Yang, J. et al. Targeting LOXL2 for cardiac interstitial fibrosis and heart failure treatment. Nat. Commun. 7, 13710 (2016).

32. Li, D., Wu, J., Bai, Y., Zhao, X. & Liu, L. Isolation and culture of adult mouse cardiomyocytes for cell signaling and in vitro cardiac hypertrophy. J. Vis. Exp. (2014). doi:10.3791/51357

33. Love, M. I., Huber, W. & Anders, S. Moderated estimation of fold change and dispersion for RNA-seq data with DESeq2. Genome Biol. 15, 550 (2014).

34. Becht, E. et al. Dimensionality reduction for visualizing single-cell data using UMAP. Nat. Biotechnol. (2018). doi:10.1038/nbt.4314

35. Hu, K. H. et al. ZipSeq: Barcoding for Real-time Mapping of Single Cell Transcriptomes. BioRxiv (2020). doi:10.1101/2020.02.04.932988

36. Stone, N. R. et al. Context-Specific Transcription Factor Functions Regulate Epigenomic and Transcriptional Dynamics during Cardiac Reprogramming. Cell Stem Cell 25, 87–102.e9 (2019).

37. Acharya, A., Baek, S. T., Banfi, S., Eskiocak, B. & Tallquist, M. D. Efficient inducible Cre-mediated recombination in Tcf21 cell lineages in the heart and kidney. Genesis 49, 870–877 (2011).

38. Kwak, H., Fuda, N. J., Core, L. J. & Lis, J. T. Precise maps of RNA polymerase reveal how promoters direct initiation and pausing. Science 339, 950–953 (2013).

39. Langmead, B. & Salzberg, S. L. Fast gapped-read alignment with Bowtie 2. Nat. Methods 9, 357–359 (2012).

40. Guo, Y., Mahony, S. & Gifford, D. K. High resolution genome wide binding event finding and motif discovery reveals transcription factor spatial binding constraints. PLoS Comput. Biol. 8, e1002638 (2012).

41. Liao, Y., Smyth, G. K. & Shi, W. featureCounts: an efficient general purpose program for assigning sequence reads to genomic features. Bioinformatics 30, 923–930 (2014).

42. Quinlan, A. R. & Hall, I. M. BEDTools: a flexible suite of utilities for comparing genomic features. Bioinformatics 26, 841–842 (2010).

43. Stadhouders, R. et al. Multiplexed chromosome conformation capture sequencing for rapid genome-scale high-resolution detection of long-range chromatin interactions. Nat. Protoc. 8, 509–524 (2013).

44. Krijger, P. H. L., Geeven, G., Bianchi, V., Hilvering, C. R. E. & de Laat, W. 4C-seq from beginning to end: A detailed protocol for sample preparation and data analysis. Methods 170, 17–32 (2020).

45. Travers, J. G. et al. Pharmacological and Activated Fibroblast Targeting of Gβγ-GRK2 After Myocardial Ischemia Attenuates Heart Failure Progression. J. Am. Coll. Cardiol. 70, 958–971 (2017).

46. Chen, E. Y. et al. Enrichr: interactive and collaborative HTML5 gene list enrichment analysis tool. BMC Bioinformatics 14, 128 (2013).

